# Decoding a cryptic mechanism of metronidazole resistance among globally disseminated fluoroquinolone-resistant *Clostridioides difficile*

**DOI:** 10.1101/2022.09.23.509282

**Authors:** Abiola O. Olaitan, Chetna Dureja, Madison A. Youngblom, Madeline A. Topf, Wan-Jou Shen, Anne J. Gonzales-Luna, Aditi Deshpande, Kirk E. Hevener, Jane Freeman, Mark H. Wilcox, Kelli L. Palmer, Kevin W. Garey, Caitlin S. Pepperell, Julian G. Hurdle

## Abstract

Severe outbreaks and deaths have been linked to the emergence and global spread of fluoroquinolone-resistant *Clostridioides difficile* over the past two decades. At the same time, metronidazole, a nitro-containing antibiotic, has shown decreasing clinical efficacy in treating *C. difficile* infection (CDI). Most metronidazole-resistant *C. difficile* exhibit an unusual resistance phenotype that can only be detected in susceptibility tests utilizing molecularly intact heme. Here we describe the mechanism underlying this trait, which we discovered using molecular genetics, phylogenetics, and population analyses. Most metronidazole-resistant strains evolved a T to G mutation, we term P*nimB^G^*, in the -10 regulatory promoter of the 5-nitroimidazole reductase *nimB*, resulting in the gene being constitutively transcribed. Silencing or deleting *nimB* eliminated metronidazole resistance. We identified the protein as a heme-dependent nitroreductase that degraded nitro-drugs to an amine lacking antimicrobial activity. We further discovered that the metronidazole-resistant P*nimB^G^*mutation was strongly associated with the Thr82Ile substitution conferring fluoroquinolone resistance in epidemic strains. Re-analysis of published genomes from global isolates confirmed that all but one encoding P*nimB^G^* also carried the Thr82Ile mutation. Our findings suggest that fluoroquinolone and metronidazole resistance co-mediated the pandemic of healthcare-associated *C. difficile* that are associated with poorer treatment outcomes in CDI patients receiving metronidazole.

## Introduction

*Clostridioides difficile* infection (CDI), a leading cause of hospital-associated diarrhea, has attracted international attention due to worsening clinical outcomes resulting from the global spread of epidemic strains of PCR ribotype (RT) 027^1–3^; RT027 belongs to phylogenetic clade 2 (Supplementary Table 1 shows the relationship between ribotype and phylogenetic classifications)^4, 5^. These strains caused outbreaks across North America, UK, and Europe with an increased incidence of severe CDI, morbidity and mortality^1, 2^. These global outbreaks also demarcate the epidemic era of CDI^3^. During this era metronidazole and vancomycin were the main two antibiotics used to treat CDI, until fidaxomicin was FDA approved in 2011. However, metronidazole is no longer recommended as a first-line drug for adult CDI, according to updated IDSA/SHEA and ESCMID CDI guidelines^6, 7^, unless fidaxomicin or vancomycin are not available. This represents a significant change in the role of metronidazole from past guidelines for CDI management^8–11^, although intravenous metronidazole combined with vancomycin is still recommended for fulminant CDI^6, 7^. These changes in CDI therapeutics warrant elucidation of the microbial genetic factors affecting efficacy of metronidazole and global CDI epidemiology. The microbial genetic determinants of metronidazole resistance are poorly understood.

Metronidazole was established as an antibiotic for CDI following clinical trials in the 1980s and 1990s, where it not only showed comparable clinical success rates to vancomycin but was much less expensive^12, 13^. However, over the last two decades it has become less effective, compared to vancomycin^14, 15^. This was originally seen in a randomized clinical trial, conducted between 1994 to 2002, where metronidazole demonstrated an 84% cure rate compared to 97% seen with vancomycin^14^. In a second clinical study conducted between 2005 to 2007, vancomycin exhibited superior cure rates to metronidazole, 81.1% compared to 72.7%^15^. These two studies reveal that metronidazole became less effective in the epidemic era. Indeed, metronidazole treatment failures in Quebec more than doubled from 9.6% in 1991-2002 to 25.7% in 2003-2004^16^, which is also the region that reported the first outbreak of epidemic RT027^2^. The reasons for the reduced clinical utility of metronidazole have been a longstanding mystery. One possibility is that increased use of metronidazole in response to rising rates of CDI imposed selection pressures that enabled the emergence and dissemination of drug-resistant *C. difficile*.

A high copy number plasmid (pCD-METRO) was recently reported to confer clinical resistance to metronidazole, in a *C. difficile* isolate obtained from a patient failing metronidazole therapy^17^. However, the overwhelming majority of metronidazole-resistant *C. difficile* do not encode pCD-METRO^17^. A survey of 10,330 publicly available genomes found pCD-METRO in just 15 strains, indicating it is exceedingly rare^18^. Further complexity arises from reports of metronidazole-resistant *C. difficile* with unstable minimum inhibitory concentrations (MICs). This has likely resulted in systematic underestimation of the prevalence of metronidazole-resistant *C. difficile*. Heme was recently shown to be essential for the accurate detection of metronidazole resistance in *C. difficile*, and the majority of metronidazole-resistant *C. difficile* exhibit heme-dependent resistance via an unknown mechanism^19, 20^; pCD-METRO does not confer heme-dependent resistance^19^. Due to the discovery of the heme-dependent phenotype, the underlying mechanism of metronidazole resistance, which was previously thought to be unstable, can now be elucidated.

Metronidazole is a nitroimidazole prodrug that is activated by cellular reduction to anion and nitroso intermediates that reacts with DNA and proteins and depletes cellular thiols^21^. Herein, we describe the discovery and genetic validation of the mechanism for heme-dependent resistance to metronidazole, followed by revealing its role in the global transmission of epidemic *C. difficile* through genome-wide association studies and population genetics. We discovered that epidemic strains evolved a common variant in the regulatory promoter of 5-nitroimidazole reductase (*CDR20291_1308*, *CdnimB*), causing its conversion from a cryptic to a constitutively expressed resistance gene. We further show that *Cd*NimB is a heme-binding flavoenzyme that bioreductively inactivates nitroimidazoles. Finally, we discovered a strong correlation between the *nimB*-promoter variant and a Thr82Ile mutation in DNA Gyrase A (GyrA), which confers fluoroquinolone resistance and has been linked to the global spread of epidemic *C. difficile*^22^. Both variations occur almost exclusively together in epidemic strains, indicating a high degree of co-selection. Therefore, we decoded cryptic mechanisms behind the rapid global spread and poor clinical outcomes of epidemic *C. difficile* strains, revealing how they were shaped by resistances to metronidazole and fluoroquinolones.

## RESULTS

### Heme mediates metronidazole resistance

We performed metronidazole susceptibility testing, with and without heme, on *C. difficile* strains of varying ribotypes and Clades (Supplementary Table 1) from the US and Europe (n=405). Forty one percent of the strains displayed a ≥ 4-fold increase in metronidazole MICs in the presence of heme (MICs=1-16 µg/ml), compared to those in the absence of heme (MICs=0.25-1 µg/ml) (Fig. 1a, Supplementary Table 1). We did not detect any metronidazole-resistant isolates that were independent of heme. These strains are designated as showing heme-dependent metronidazole resistance. Conversely, metronidazole-susceptible strains displayed a 2-fold or less difference in MICs, with or without heme.

**Figure 1.**
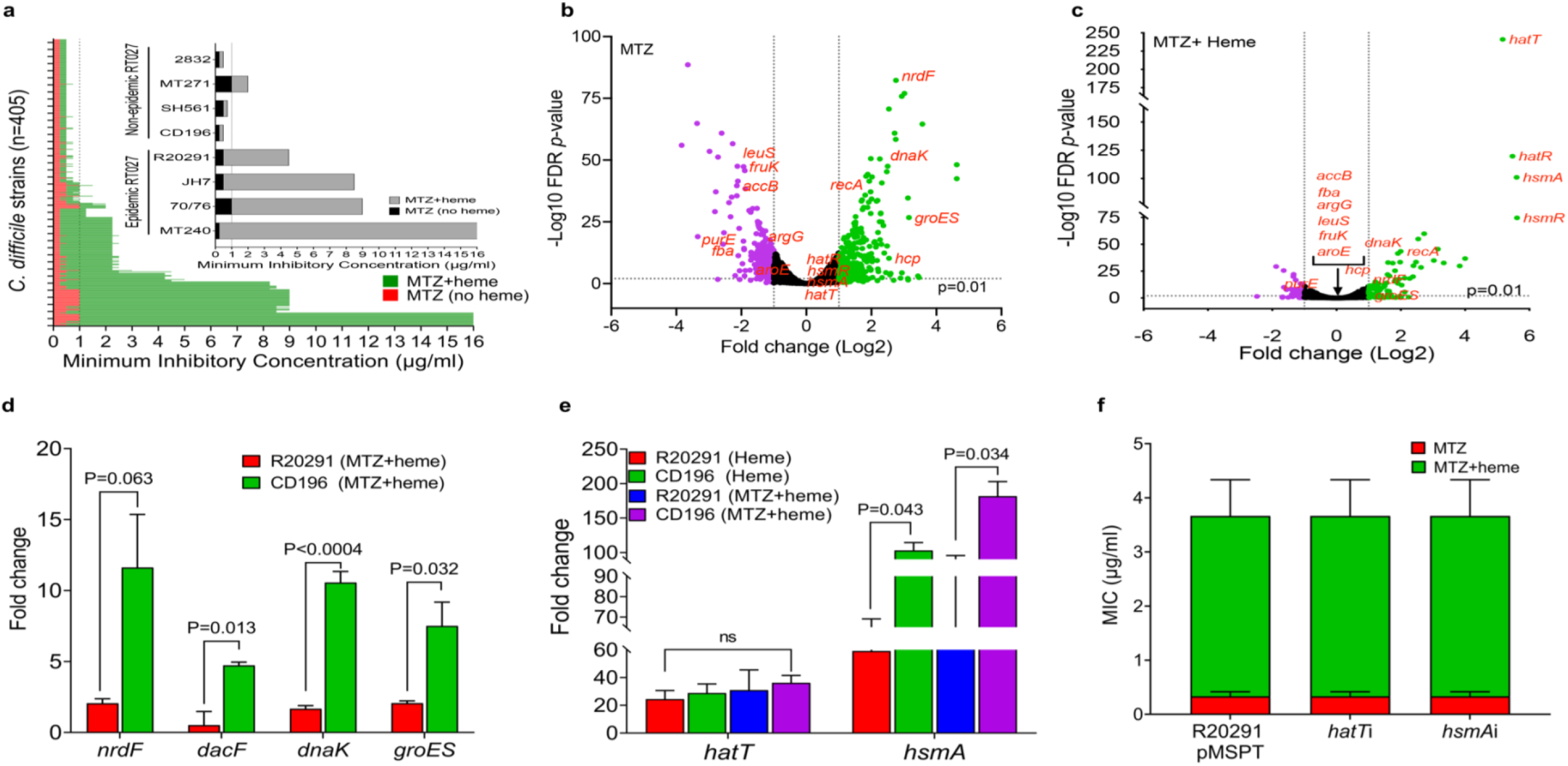
Heme attenuates cellular toxicity of metronidazole (MTZ) in epidemic *C. difficile* strains. **(a)** Minimum inhibitory concentrations (MICs) of MTZ were determined against global clinical isolates of *C. difficile* (n=405) in the presence and absence of heme. Isolates with heme-dependent MTZ resistance exhibited ≥4-fold higher MICs in BHI agars containing heme, compared to agars without heme. The line at 1 μg/ml indicates MTZ MIC found to associated with poor clinical outcomes^44^ (the EUCAST breakpoint is 2 μg/ml). Inset illustrates selected RT027 epidemic and non-epidemic strains that are resistant or susceptible to MTZ in heme, respectively. Non-epidemic RT027 were defined as previously described^22^ (lacking fluoroquinolone resistance SNPs in gyrase A and large transposons found in epidemic RT027). MICs were from two biological replicates with two technical replicates. **(b, c)** Transcriptome response of epidemic R20291 (OD600nm ≈ 0.2) exposed for 30 min to MTZ (2 μg/ml) in the absence (**b**) and presence (**c**) of heme (5 μg/ml). The volcano plots indicate differentially expressed genes and their statistical significance; the purple and green dots indicate significantly downregulated and upregulated genes, respectively; Red-highlighted genes are cell stress-responsive and metabolic genes whose expression were altered by MTZ but were attenuated by heme. The RNAseq is based on two biological replicates, as the third replicate was rejected in FastQC due to poor quality. Pearson correlation of R^2^= 0.94 in Extended Fig. 1b shows RNAseq transcriptional changes were validated by qPCR (three biological replicates). **(d)** Transcriptional response of epidemic R20291 and non-epidemic CD196 to MTZ (2 μg/ml) in the presence of heme (5 μg/ml). qPCR analysis of transcription patterns of genes indicative of MTZ toxicity demonstrates heme is not protective for non-epidemic CD196; statistical significance was assessed by unpaired t-test from three biological replicates in Grapdpad prism 9.4.1. **(e)** Transcriptional analysis of heme sensing/detoxifying genes *hatT* and *hsmA* in heme with or without MTZ, indicate that both CD196 and R20291 both highly express these genes, as determined by qPCR; statistical significance was assessed by unpaired t-test from three biological replicates. **(f)** Heme sensing and detoxifying systems do not mediate heme-dependent MTZ resistance. In R20291, silencing *hatT* or *hsmA* had no effect on heme-dependent resistance, indicating that these genes are unlikely to contribute to heme-dependent resistance; anti-sense RNA to *hatT* or *hsmA* was cloned into vector pMSPT and induced by 64 ng/ml of anhydrotetracycline; qPCR indicated *hatT* mRNA decreased by ∼7-fold (i.e., -7.21 ± 1.95 of 3 replicates) and *hsmA* by ∼18-fold (i.e., -18.46 ± 10.66 of 3 replicates). Data in **d** to **f** are plotted as mean ± standard error of mean.

### Transcriptome analysis confirms heme attenuates cellular toxicity of metronidazole

R20291, a well-known model strain of epidemic RT027 isolated from a hospital outbreak in 2003-2006 in Stoke Mandeville, UK^23^, was adopted as our main research strain since it shows heme-dependent resistance to metronidazole (MICs of 2-4 µg/ml and 0.25-0.5 µg/ml, with and without heme, respectively). Examination of transcriptional responses of R20291 exposed to metronidazole (2 µg/ml) and heme (5 µg/ml) for 30 mins showed that heme attenuated the upregulation of pathways associated with metronidazole toxicity. Without heme, there were 285 upregulated genes (≥2-fold) by metronidazole, but only 44 of these genes were upregulated when heme was present (Fig. 1b-c). Pathways induced by metronidazole toxicity that were mitigated by heme include DNA replication and repair (e.g., ribonucleotide reductase, *nrdF*) and stress responsive chaperones (e.g., co-chaperonin, *groES* and bacterial Hsp70 chaperone, *dnaK*). By itself, heme minimally affected *C. difficile* transcriptome (Extended Data Fig. 1a). Validation of the overall quality of the RNA-seq data by qRT-PCR showed a high correlation of R^2^= 0.94 between both datasets (Extended Data Fig. 1b-c). We next tested whether heme could also quench metronidazole toxicity in metronidazole-susceptible *C. difficile* CD196 (a pre-epidemic RT027 strain that was isolated in 1985 in Paris). However, heme did not attenuate gene expression associated with metronidazole toxicity in CD196 (Fig. 1d, Extended Data Fig. 1d-e). It was postulated that the heme-binding transcriptional regulator HsmA^24^ may be associated with heme-dependent metronidazole resistance, as the gene appears inactivated in RT010 strains^19^. However, we did not find any evidence for the involvement of HsmA or the heme detoxifying system encoded by *hatRT* ^25^ in mediating heme-dependent metronidazole resistance (Fig. 1e-f). Collectively, the data suggested that other genetic factors mediate heme-dependent metronidazole resistance.

### Identification of 5-nitroimidazole reductase (NimB) as a mechanism for *C. difficile* heme-dependent metronidazole resistance

To screen for genetic loss of heme-dependent metronidazole resistance we generated ∼7,488 transposon (Tn) mutants in R20291 using vector pRPF-215 that carries an ATc-inducible *Himar1*^26^. This identified two Tn mutants that were 8-fold more susceptible to metronidazole (MICs of 0.5 µg/ml), compared to the wildtype (MIC of 4 µg/ml). Genome sequencing revealed that the two Tn mutants each had single insertions in *CDR20291_1308* or *CDR20291_2676* at positions 1547478 (CDS position 464 of 468 nt) and 3152802 (CDS position 1151 of 2412 nt), respectively in the R20291 genome (FN545816.1).

The two genes are annotated as 5-nitroimidazole reductase (*nimB*) and cysteine protease (*cwp84*), respectively. Cwp84 is responsible for cleaving the surface layer (S-layer) protein precursor SlpA into low- and high-molecular weight subunits that form the paracrystalline S-layer. Although *cwp*84 is a non-essential gene, deletion mutants have slower growth rates, which might impose pleiotropic effects that increase metronidazole susceptibility of the *cwp84* Tn mutant^27^. *C. difficile* NimB (*Cd*NimB) belongs to the pyridoxamine 5’-phosphate oxidase family of proteins and is closely related NimAs from *Terrisporobacter* spp. (60% identity) and *Bacteroides fragilis* (45% identity). Nim proteins are thought to confer resistance to nitromidazoles by converting their nitro group to antimicrobial inactive amino derivatives, hence avoiding the generation of reactive species^28^. It is unclear, however, to what extent *nim* genes confer resistance to metronidazole since their presence does not directly result in resistance.

Despite this, we chose to further study *nimB* because of the marked phenotype of the *Tn* mutant (R20291-*Tn::nimB*) and the high likelihood that in *C. difficile nimB* may be another cryptic resistance mechanism^29^. Hence, when we generated a *nimB* knockout mutant by allelic exchange in R20291, the mutant recapitulated the metronidazole-susceptible phenotype of R20291-*Tn::nimB* (MIC=0.25 µg/ml vs. 4-8 µg/ml for the wildtype in the presence of heme). Importantly, heme-dependent resistance was restored (MICs=4-8 µg/ml) upon complementation of R20291-*Tn::nimB* and R20291Δ*nimB* with wildtype *nimB* from R20291 (Fig. 2a).

**Figure. 2.**
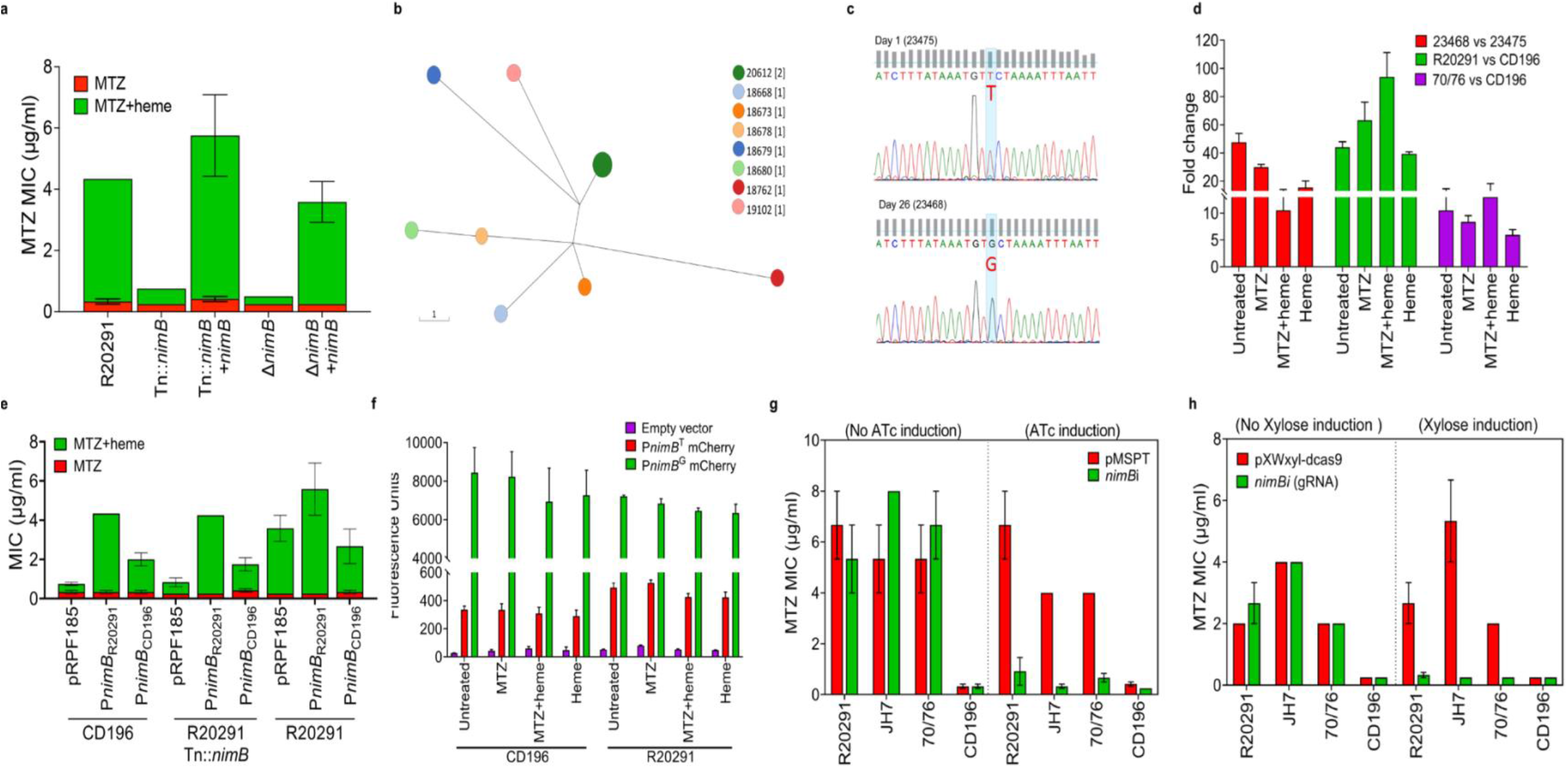
Constitutive transcription of *nimB* is associated with metronidazole (MTZ) resistance and results from a mutation in the -10 promoter. **(a)** Screening of a transposon (Tn) library identified that insertional inactivation of *nimB* abrogated resistance in R20291. The allelic deletion of *nimB* confirmed its role in resistance; in both mutants, resistance was restored by complementation with wildtype *nimB*, expressed from its native promoter. **(b)** Sequential isolates from a patient failing MTZ therapy were collected on day 1 (before therapy) and on day 26 (after therapy ended); the day 1 isolate (23475, MIC=0.25 μg/ml) and day 26 isolate (23468, MIC=1.0 μg/ml) were MTZ-susceptible and -resistant, respectively. In Enterobase, core-genome multilocus sequence typing showed the two isolates had the same hierarchical clustering (HC0), indicating they have identical core genomes or at least do not differ by more than two SNPs. cgMLST Ninja Neighbor Joining GrapeTree of *C. difficile* belonging to ST350; 23475 and 23468 clustered in the same node. The scale bar indicates the number of allelic differences. **(c)** Sanger sequencing showed 23468 has a T to G mutation in the -10 *nimB*-promoter when compared to 23475, confirming a whole genome comparison; this mutation is referred to as P*nimB^G^* in 23468, while the wildtype is designated P*nimB^T^* in 23475. **(d)** qPCR analysis of *nimB* transcription in resistant and susceptible strains carrying P*nimB^G^* and P*nimB^T^*, respectively. Resistant strains (23468, R20291, and 70/76) had higher levels of *nimB* mRNA than susceptible strains (23475 and CD196); *nimB* transcription was constitutive in resistant strains. Strains were cultured in various conditions (MTZ [2 μg/ml] and heme [5 μg/ml]); shown are fold mRNA amounts in resistant strains relative to susceptible strains. **(e)** Relationship between resistance and variations in *nimB* between non-epidemic CD196 and epidemic R20291. The *nimBs* of CD196 and R20291 were expressed from their native promoters in the indicated strains. **(f)** Comparison of the promoter strengths of P*nimB^G^*and P*nimB*^T^ based on the transcription and fluorescence of mCherryOpt. Fluorescence was normalized to OD600nm culture density. P*nimB^G^*was associated with constitutive expression, reflected by higher fluorescence. **(g, h)** Genetic silencing of *nimB* reverses resistance. *nimB* was silenced by an antisense RNA (asRNA) (**g)** and by CRISPR interference with two guide RNAs (gRNAs) (**h**). AsRNA was induced by anhydrotetracycline (64 ng/ml) from P*tet* promoter in vector pMSPT, while gRNA was induced by xylose (1% w/v) from P*xyl* promoter in vector pXWxyl-dcas9. Data are shown as the mean ± standard error of mean from three biological replicates.

### Heme-dependent resistance results from a SNP in *nimB* -10 promoter

To identify genetic mechanisms in clinical strains we sequenced a pair of isolates obtained from a single patient, before and after therapy with metronidazole; these isolates were from the MODIFY clinical trials of bezlotoxumab^30^. The metronidazole-susceptible isolate (23475) was recovered at baseline diagnosis and had an MIC of 0.25 µg/ml, while the second isolate (23468) obtained after metronidazole therapy (day 26 after diagnosis) had an MIC of 1 µg/ml (no other patient information was disclosed). To assess genetic relatedness, the genomes of 23475 and 23468 were analyzed by cgMLST in EnteroBase, which indicated that they formed a unique HC0 hierarchical cluster that is indicative of an indistinguishable core-genome sequence type (Fig. 2b). Genome alignment and Sanger sequencing revealed that the resistant strain 23468 evolved a single nucleotide polymorphism (SNP) of T to G in the predicted -10 regulatory promoter region (i.e., TTCTAAAAT to TGCTAAAAT) (Fig. 2c). The SNP differences were designated as P*nimB^T^* and P*nimB*^G^, for the susceptible wildtype and resistant mutant, respectively. Because the -10 region influences RNA polymerase activity^31, 32^, we tested whether the SNP affected *nimB* transcription. qRT-PCR revealed 23468 had on average 16- to 40-fold greater amounts of *nimB* transcripts compared to 23475 (Fig. 2d). Elevated *nimB* transcription in 23468 was not dependent on heme or the presence of metronidazole, suggesting the gene was constitutively transcribed. Constitutive expression may also explain why differential expression of *nimB* was not seen in our RNAseq. We next compared the regulatory regions of *nimB* in pre-epidemic CD196 (susceptible) with epidemic R20291 (resistant). This revealed that CD196 encoded P*nimB^T^*, whereas R20291 encoded P*nimB^G^*, suggesting that evolution of a SNP in the regulatory region of *nimB* engendered metronidazole resistance. We therefore compared *nimB* mRNA levels in different metronidazole-susceptible strains (i.e., CD196 and CD630 carrying P*nimB*^T^) and resistant strains (i.e., R20291 and 70/76 carrying P*nimB^G^*). R20291 exhibited ∼39±1- to 94±17-fold or 81±5- to 203±100-fold greater *nimB* mRNA relative to the susceptible strains (CD196 and CD630), with or without heme or metronidazole (Fig. 2d, Extended Data Fig. 2a). The resistant strain 70/76 also had greater amounts of *nimB* mRNA, compared to susceptible strains (Fig. 2d, Extended Data Fig. 2a). Overall, all resistant strains had lower CT values, indicative of elevated *nimB* mRNA when compared to susceptible strains (Extended Data Fig. 2b).

### P*nimB^G^* enhances transcription of *nimB*

Gene expression from P*nimB^T^* and P*nimB^G^*was examined in several ways to establish why *nimB* mRNA levels differ. Firstly, *nimB* from CD196 or R20291 were ectopically expressed under their cognate promoters in susceptible CD196 and R20291-*Tn*::*nimB*. In susceptible backgrounds, the P*nimB^T^ nimB* sequence from CD196 conferred lower MICs (1-2 μg/ml) than P*nimB^G^* from R20291 (MICs of 4-8 μg/ml) (Fig. 2e). Secondly, because NimB of CD196 and R20291 differ by a single Leu155Ile polymorphism, we examined the effect of this difference by expressing both genes under the tetracycline-inducible promoter (P*tet*). Under P*tet*, both polymorphisms conferred equivalent heme-dependent resistance in R20291-Tn::*nimB* (Extended Data Fig. 2c), contradicting a previous hypothesis that Leu155Ile influences metronidazole resistance^33^. Thirdly, we used the mCherryOpt reporter^34^ to compare the promoter strengths of P*nimB^T^* and P*nimB^G^*in different genetic backgrounds, under drug-free and metronidazole conditions. As shown in Fig. 2f, CD196 expressing P*nimB^G^*::mCherryOpt had higher fluorescence (6943±1729 to 8437±1308 RFU) when compared to P*nimB^T^*::mCherryOpt (290±43 to 335 ±26 RFU). Similarly, R20291 expressing P*nimB^G^*::mCherryOpt displayed higher fluorescence than P*nimB^T^*::mCherryOpt (6354±454 to 7219±56 RFU and 423±38 to 527±20 RFU, respectively). These trends were consistent in other susceptible and resistant strains tested i.e., P*nimB^G^*::mCherryOpt ranged from 2932±508 to 5222±492 RFU and P*nimB^T^*::mCherryOpt from 268±18 to 436±11 RFU (Extended Data Fig. 2d-e); qPCR confirmed that the higher mCherry fluorescence from P*nimB^G^* corresponded with greater amounts of mCherry mRNA, compared to P*nimB^T^* (Extended Data Fig. 2f). Fourthly, since *nimB* is constitutively transcribed under P*nimB^G^*, we reasoned genetic silencing would restore susceptibility to metronidazole. We therefore silenced *nimB* with two different methods^35^, with vector pMSPT, considered to block mRNA translation by antisense RNA (asRNA) targeting the UTR region, or by Crispr-interference to block transcription with a guide RNA (gRNA) to *nimB* expressed from vector pXWxyl-dcas9^35^. Expression of asRNA to *nimB* from pMSPT reversed heme-dependent resistance in R20291 and other RT027 strains tested, compared to the empty vector or uninduced controls (Fig. 2g). Silencing *nimB* with gRNA also increased the susceptibility of these strains to metronidazole, when compared to the empty vector or uninduced controls (Fig. 2h). Taken together, these data show heme-dependent metronidazole resistance involved constitutive transcription of *nimB* and P*nimB^G^* is a stronger promoter.

#### *C. difficile* NimB (*Cd*NimB) is a heme-binding protein

We next revealed that NimB is a hemoprotein, explaining why heme is required for metronidazole resistance. There are no reports that Nim proteins bind heme. However, Phyre2-predicted the *Cd*NimB was structurally related to the heme binding flavocytochromes Anf3 nitrogenase from *Azotobacter vinelandii* (PDB ID 6RK0) and MSMEG_4975 flavin/deazaflavin oxidoreductases from *Mycobacterium smegmatis* (PDB ID 4YBN)^36, 37^. Anf3 and MSMEG_4975 are pyridoxamine 5’-phosphate oxidase family proteins that bind heme through the homologous proximal histidine-70 and histidine-62, respectively^36, 37^. To discover the heme binding motif(s) in NimB proteins, we first aligned the *B. thetaiotaomicron* NimB crystal structure (PDB 2FG9) with that of Anf3, revealing that *Bt*NimB histidine-50 was homologous to Anf3 histidine-70 (Extended data Fig. 3). Similarly, sequence alignment and structural homology modeling indicated that histidine-55 of *Cd*NimB was homologous to Anf3 histidine-70 (Fig. 3a-b). Interestingly, the amino acid is also identical to histidine-71 in *Deinococcus radiodurans* (*Dr*NimA, Q9RW27; Fig. 3a), which was hypothesized to bind an unidentified cofactor that affected the color of the enzyme in suspension^38^.

**Figure 3.**
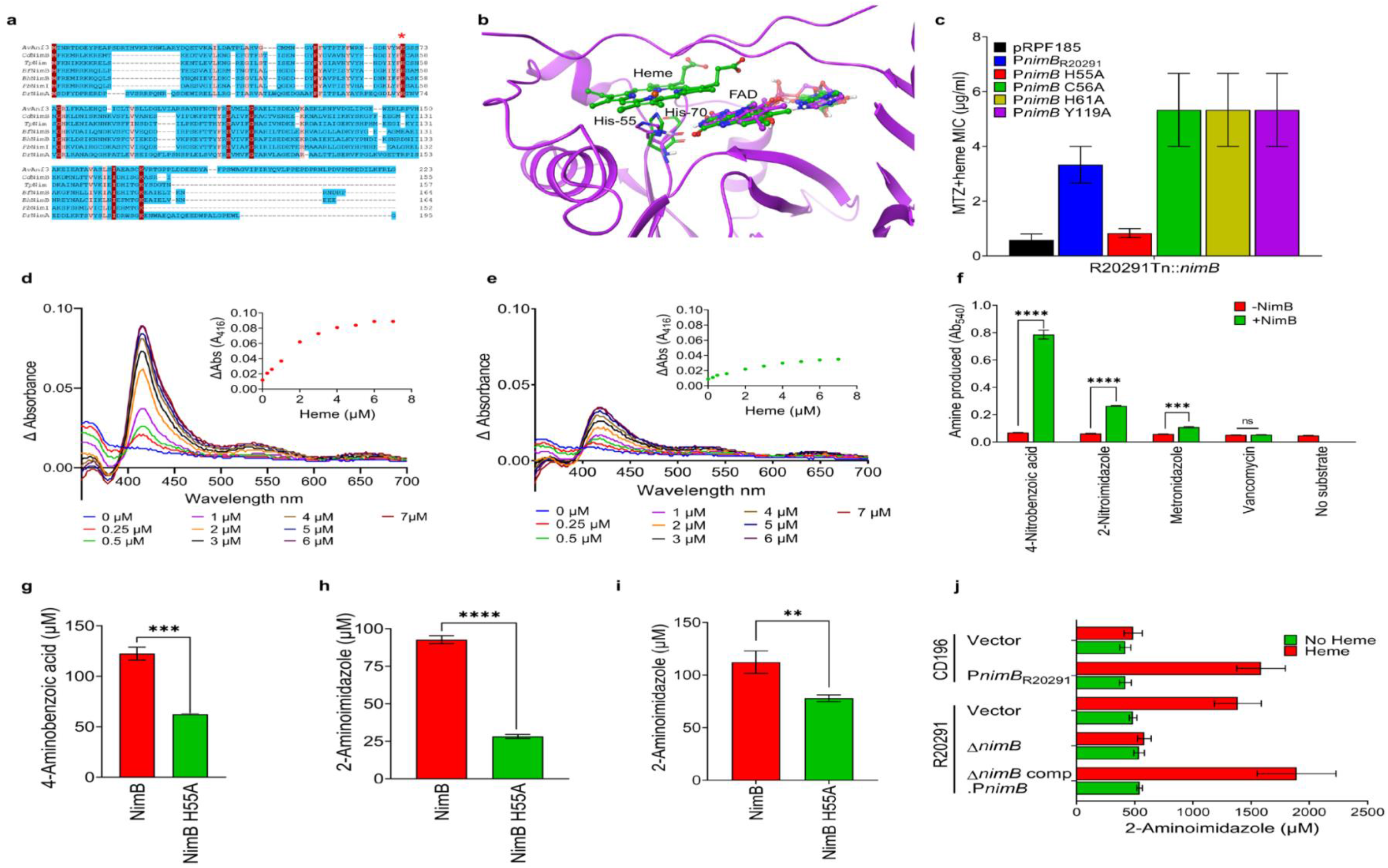
*Cd*NimB is a heme-binding nitroreductase. **(a)** Sequence alignment of Nim proteins from *C. difficile* and other bacteria, with the structurally related heme binding flavocytochrome Anf3 nitrogenase from *Azotobacter vinelandii* (AvAnf3, accession no. 6RK0). Histidine-70 (red asterix) is the proximal heme-binding ligand of Anf3 and a conserved amino acid in Nim proteins e.g., histidine-55 in *Cd*NimB. Other sequences and accession numbers are from: *Tp*, *Terrisporobacter petrolearius* WP_228108130; *Bf*, *Bacterioides fragilis* WP_063854490.1; *Bh, Brachyspira hampsonii* WP_039955657; *Dr*, *Deinococcus radiodurans* Q9RW27; and *Pb*, *Prevotella baroniae* ACR40098.1 **(b)** Structural alignment with Anf3 (green ball and stick), a *Cd*NimB homology model (purple ribbons) shows a close structural similarity. The *Cd*NimB model was generated from the *B. thetaiotaomicron* NimB X-ray structure bound to FAD (PDB 2FG9) and aligned with Anf3, which has both FAD and heme bound (see Extended Fig. 3). The model shows *Cd*NimB histidine-55 (purple) is structurally equivalent to histidine-70 (green) of Anf3; the FAD domains are also conserved in the two proteins. FAD and heme of Anf3 are represented as ball/stick with green carbons; the FAD of *Cd*NimB is shown as ball/stick with purple carbons; for simplicity only histidine-70 of the Anf3 protein is shown. **(c)** Alanine mutagenesis identified histidine-55 mediates heme-dependent metronidazole resistance (MTZ). Among different alanine mutants, only His55Ala mutant did not exhibit resistance when expressed in R20291-*Tn::nimB*; the *nimB* variants were expressed in pRPF185 under P*nimB^G^*. (**d, e)** Absorption spectra showing wildtype *Cd*NimB (**d**) binds heme, but this is attenuated in the His55Ala *Cd*NimB mutant (**e**). The spectra were obtained by adding increasing concentrations of heme (i.e., hemin, 1-7 µM) to 10µM of protein (data is representative of three experimental replicates). Heme was titrated until saturation was reached, where there were no further significant changes in absorbance readings. The buffer was similarly titrated with heme and values subtracted from the protein spectral data. The insets show binding saturation curves based on the change in absorbance at 416 nm as a function of heme concentration. **(f)** Nitroreductase activity of wildtype *Cd*NimB. Reduction of various nitroaromatics were tested in reaction containing 10 μM of *Cd*NimB and 10 μM of heme. Reactions were incubated for 2 h, and formation of aromatic amines detected using Bratton-Marshall assay; vancomycin, a non-nitroaromatic was a negative control. **(g, f)** Comparison of nitroreductase activities of wildtype *Cd*NimB and His55Ala mutant, with quantification using the Bratton-Marshall assay. As shown in **g**, 4-nitrobenzoic acid is reduced to 4-aminobenzoic acid, and 2-nitroimidazole is reduced to 2-aminoimidazole in f. **(h)** LC-MS/MS quantification of 2-aminoimidazole formed from the reduction of 2-nitroimidazole in nitroreductase assays with wildtype or His55Ala *Cd*NimBs. **(i)** Cellular reduction of 2-nitroimidazole to 2-aminoimidazole. Concentrated cultures of various isogenic strains were treated with 2-nitroimidazole (2 mM) alone or with heme and incubated for 3 h, before 2-aminoimidazole were quantified. Throughout, data are from at least three biological replicates; plots show the mean ± standard error of mean (****P≤0.0001, ***P≤0.001, **P≤0.01, and ns= not significant).

Hemoquest^39^, an algorithm that predicts heme-binding motifs, also identified *Cd*NimB histidine-55 as a candidate heme binding residue, along with cysteine-56, histidine-61, cysteine-101, and tyrosine-119. Therefore, to define the crucial residue, NimB variants (His55Ala, Cys56Ala, His61Ala, and Tyr119Ala) were created by site-directed mutagenesis, expressed in susceptible R20291-*Tn::nimB* and tested for loss of resistance. The Ala-55 mutant was the only variation that did not cause heme-dependent resistance (Fig. 3c, Extended Data Fig. 4a).

**Figure 4.**
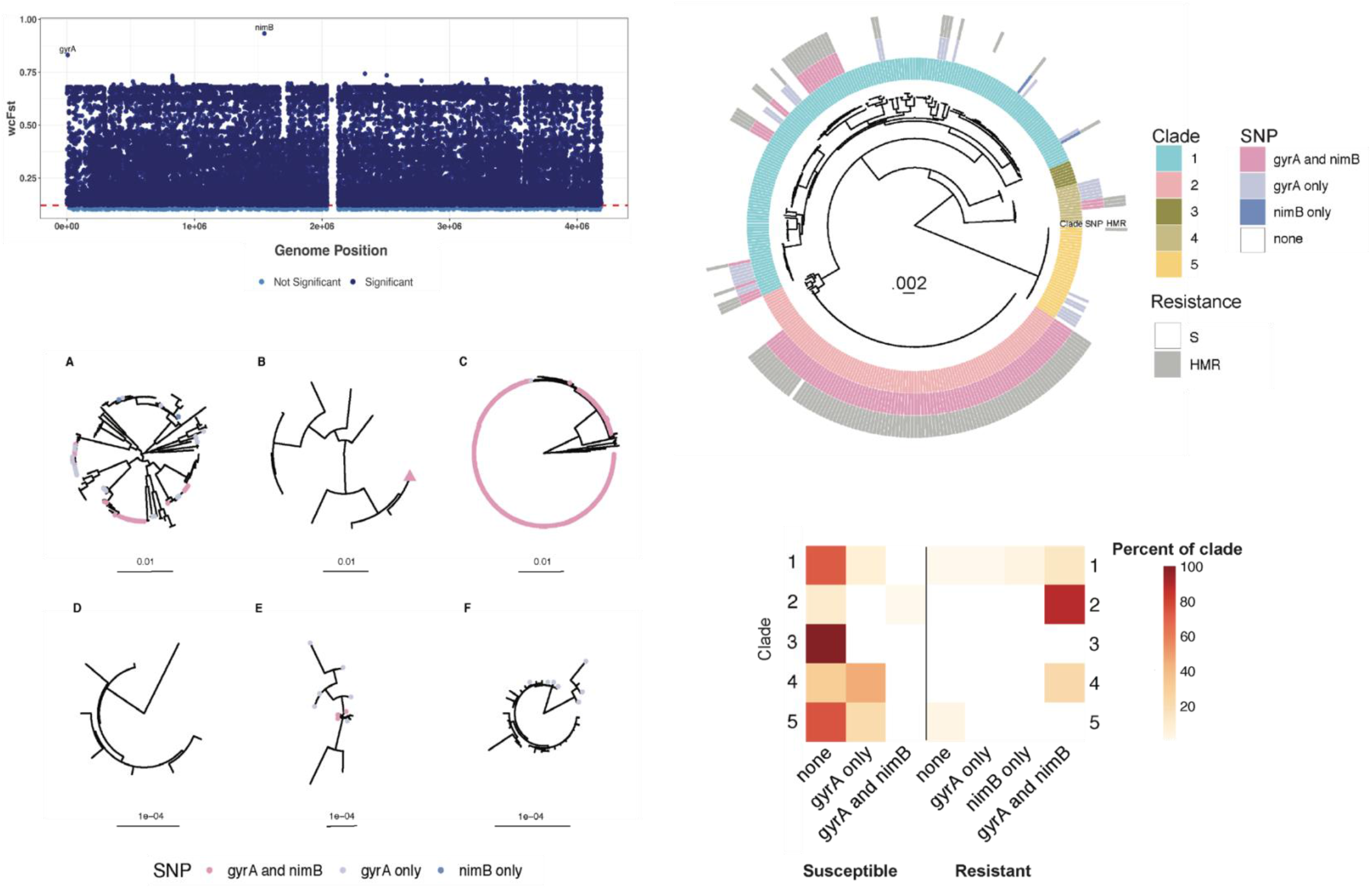
P*nimB^G^*-associated metronidazole (MTZ) resistance co-occurs with fluroquinolone resistance and pandemic spread. **(a)** Weir & Cockerham’s Fixation index (FST) was calculated for all SNPs, comparing heme-dependent metronidazole-resistant (HMR) and susceptible (S) populations. FST values are plotted by genome position. Both *gyrA* and *nimB* SNPs showed exceptionally high FST values, indicating extreme frequency differences between HMR and S populations. (**b**) Maximum likelihood phylogeny based on whole genome sequences from 348 isolates. Multi-locus sequence type (MLST) clades 1-5 (inner circle), *gyrA* and *nimB*-promoter mutations (middle circle), and metronidazole susceptibility (outer circle), either heme-induced metronidazole resistant (HMR) or metronidazole susceptible (S), are shown. The *gyrA*, *nimB*-promoter mutations and associated HMR are associated with clonal groups in Clades 2 and 1, suggesting that bacteria carrying these variants have spread rapidly and/ or are sampled more densely in this population. (**c**) Maximum-likelihood phylogenies of individual clades 1-5 with tips colored according to *gyrA* and *nimB* SNP presence or co-occurrence from this study as well as He et al.^22^ Shown are: A, Clade 1; B, Clade 2, including ‘global’ isolates from He et al.^22^ and those in this study. The pink triangle represents the clonal group of which all but one have both SNPs; C, Inset showing detail of the clonal group within Clade 2 (with one exception (blue dot; *gyrA* only), isolates with the *gyrA* mutation also encoded the *nimB* mutation; D, Clade 3; E, Clade 4; and F, Clade 5. (**d**) Heat map showing SNP presence/co-occurrence and MTZ resistance phenotype frequencies within each clade. Each row corresponds to a single clade. Each column presents a summary of *gyrA* and *nimB* SNP presence and MTZ resistance phenotype. Heat map is shaded by the percentage of each condition within each clade, such that the sum across a single row is 100%. 88% of Clade 2 isolates are HMR with both *gyrA* and *nimB* SNPs, while 100% of Clade 3 isolates are MTZ susceptible and have neither SNP of interest.

Next, we directly compared heme binding by wildtype *Cd*NimB and the NimB-Ala-55 proteins. Incubation of wildtype *Cd*NimB (10 μM) from R20291 with increasing amounts of heme (1 to 7 μM) produced a Soret band at 416 nm (Fig. 3d), confirming the development of a heme-protein complex. Increasing concentrations of heme caused higher peak intensity, reaching saturation at ≥4 μM of heme (Fig. 3d). In contrast, equimolar NimB-Ala-55 displayed at least 50% lower Soret band intensities (Fig. 3e), indicating reduced formation of the protein-heme complex. This was confirmed visually, as enzyme-heme suspensions from the wildtype *Cd*NimB-heme suspensions were more intensely reddish-brown color, indicating heme-protein complexation (Extended Data Fig. 4b). Even though heme was not previously recognized as a cofactor for Nim proteins, purified *Dr*NimA showed a Soret band, which is absent in His-71 mutants^38^. Pyruvate was previously thought to be a cofactor^40^, but this was also disproved^38^. Consequently, *Cd*NimB is a heme-binding protein, with histidine-55 most likely acting as its proximal ligand, a characteristic expected to be conserved in other Nim proteins.

### NimB inactivates nitroaromatic compounds in a heme-dependent manner

Mechanistically, Nim proteins are thought to transfer two electrons from an unknown cofactor to convert nitro-groups into non-toxic amino derivatives^28, 38, 41^. In support of Nim inactivating nitroimidazoles, *B. fragilis* NimA cells formed dimetridazole amino derivatives in Wilkins-Chalgren medium containing heme^28^. To test the nitroreductase activity of *Cd*NimB under anaerobic conditions, we added heme to a nitroreductase assay^36^ in which NADPH (electron donor) oxidation is coupled to FAD reduction. The Bratton-Marshall reagent that is specific for primary aromatic amines was then used to colorimetrically detect the aromatic amino product under anaerobic conditions. As shown in Fig. 3f, *Cd*NimB converted 4-nitrobenzoic acid, metronidazole, and 2-nitroimidazole to their amino counterparts.

To quantify amino reaction end-products of *Cd*NimB enzymatic and cellular nitroreductase activities we adopted the related 2-nitromidazole, since 5-aminoimidazoles derived from 5-nitroimidazoles (e.g., metronidazole) are chemically unstable, undergoing ring cleavage in air^42^, which would present challenges to quantify by LC-MS/MS. In contrast, 2-nitroimidazole produces more stable 2-aminoimidazoles^42^ and are substrates for *Cd*NimB (Fig. 3f). *C. difficile* also showed heme-dependent resistance to 2-nitroimidazole i.e., in the absence of heme, MICs were 0.5-1, 0.25 and 0.25 µg/ml, against susceptible CD196, R20291Δ*nimB* and R20291-Tn::*nimB*, respectively but were 4-8 µg/ml against resistant R20291 (Extended Data Fig. 4c). While complementation of the susceptible strains with wildtype *nimB*, from R20291, restored resistance to 2-nitroimidazole, complementation with the Ala-55 variant did not (Extended Data Fig. 4c). In nitroreductase enzyme assays, NimB-Ala-55 was less efficient in reducing 4-nitrobenzoic acid and 2-nitroimidazole (Fig. 3g-h). LC–MS/MS analysis confirmed that 2-nitroimidazole was converted to 2-aminoimidazole by wildtype *Cd*NimB, whereas the NimB His55Ala variant made considerably less of the amino product (Fig. 3i). We next examined cellular reduction of 2-nitroimidazole, using isogenic strains of R20291 (with an empty vector control, R20291Δ*nimB*, and R20291Δ*nimB::nimB* complementation) and CD196 (with an empty vector control or ectopically expressing *nimB* from P*nimB^G^*). In the absence of heme, both susceptible and resistant strains produced close to equal, but low, amounts of 2-aminoimidazole (∼419± 48 to 542±24 μM), as measured by LC-MS/MS. Adding heme increased the production of 2-aminoimidazole by ∼3-4-fold (i.e., ∼1386±201 to 1890±340 μM), but in only the resistant strains (Fig. 3j). These findings indicate that *Cd*NimB is a nitroreductase that converts nitroimidazoles to amino derivatives enzymatically and cellularly, in a heme-dependent manner.

### Evolutionary analysis of metronidazole resistance in clinical isolates

Since heme-dependent metronidazole resistance appears to be the dominant phenotype of metronidazole-resistant *C. difficile*, we decided to further investigate potential mechanisms underlying this phenotype with genome-wide association studies (GWAS) of geographically diverse natural bacterial populations. We performed these analyses on a dataset of 348 clinical isolates for which we determined MICs of metronidazole in the presence and absence of heme. Following a previously reported approach^43^, we used fixation index (FST) outlier analysis to identify candidate genetic variants associated with heme-dependent metronidazole resistance. Among 225,113 SNPs segregating in the sample, we identified two SNPs as extreme outliers, with marked differentiation in frequency between metronidazole-susceptible and -resistant populations (Fig. 4a). The *nimB*-promoter mutation was one of these SNPs and, surprisingly, the other variant associated with resistance was a mutation 245C>T in *gyrA* (Thr82Ile in gyrase A) that confers resistance to fluoroquinolones. The Thr82Ile mutation has been associated with pandemic spread of health care associated *C. difficile*.^22^ There is no known mechanistic basis for an association between *gyrA* and metronidazole resistance. Instead, we hypothesize that co-occurrence of the Thr82Ile in GyrA and the *nimB*-promoter SNP are both strongly selected, and that their combination enabled pandemic spread of *C. difficile*. To test this hypothesis, we first investigated the genomic data for signs of selection by identifying homoplasies: variants that arise independently more than once on the phylogeny and are a potential signal of positive selection^43^. Using ancestral reconstruction, we estimated that the *gyrA* mutation arose independently on 17 occasions in our sample of 348 strains, whereas the *nimB*-promoter mutation arose 15 times, placing both variants in the 99.9^th^ percentile of homoplasies. These findings are indicative of positive selection and suggest that the use of metronidazole, in addition to fluoroquinolones, exerted important selection pressures shaping recent adaptation of *C. difficile*.

A maximum likelihood phylogeny inferred from whole genome sequence data is shown with associated metronidazole susceptibility, clade and *gyrA*/*nimB*-promoter allele indicated for each isolate in the sample (Fig. 4b). The phylogeny shows that the P*nimB^G^* mutation occurs almost exclusively in association with the *gyrA* mutation. While more isolates (n=34) carried the *gyrA* mutation alone than the *nimB* mutation (n=2), most isolates with the *gyrA* mutation also carried the P*nimB^G^*variant. *C. difficile* can be clustered into five major clades^5^. In our sample, these co-occurring variants were primarily associated with clade 2 and to a lesser extent clade 1, whereas they were absent from clades 3 and 5 (Fig. 4d).

Within clades 1 and 2, the *gyrA* and P*nimB^G^* mutations were associated with large clonal groups (Fig. 4b), suggesting that strains with these variants were rapidly transmitted. To test this hypothesis, we re-analyzed the dataset from He *et al.*^22^ who previously determined that *gyrA* mutations were associated with global epidemic spread of health care associated *C. difficile*. Our re-analysis of these published genomes showed that all but one of the isolates carrying the *gyrA* mutation also carried the P*nimB^G^* variant. Together, these observations suggest that mutations at the *nimB*-promoter site are responsible for the vast majority of metronidazole resistance (Fig. 4c) and thus likely played a pivotal role in declining efficacy of this treatment. We further link metronidazole resistance with fluroquinolone resistance and find evidence suggesting the combination of resistances underlies the recent global spread of health care associated *C. difficile* diarrhea.

Our study also identified heme-dependent metronidazole-resistant strains that lacked the P*nimB^G^* SNP. Molecular analysis of two of these strains (i.e., 17/27 and 25603) showed that their resistance phenotypes also depended on *nimB*. In both strains, *nimB* mRNA levels were higher than those in susceptible CD196 or CD630, and silencing *nimB* rendered them more susceptible to metronidazole (Extended Data Fig. 5a-d); this observation is analogous to that for genetically analyzed strains with P*nimB^G^*(Fig. 2d, 2f-g; Extended Data Fig. 2a-c). Accordingly, whereas most heme-dependent metronidazole-resistant strains encode P*nimB^G^*, a minority appear to have alternative transcriptional or post-transcriptional mechanisms to regulate *nimB*.

## Discussion

Over the last two decades, the global spread of epidemic *C. difficile* has been paralleled by declining response rates of metronidazole, culminating in major guidelines removing it as a recommended first-line option for CDI. We recently demonstrated that heme-dependent resistance to metronidazole, defined by an MIC breakpoint of ≥1 µg/mL, was associated with an increased risk of metronidazole clinical failure^44^. Fundamental to this finding was the revelation that heme is crucial for detecting metronidazole-resistant *C. difficile*^44^. Our findings that most metronidazole-resistant strains evolved a T to G mutation (P*nimB^G^*) in the -10 promoter of *nimB* provide a microbial genetic explanation as to why such strains might be less responsive to metronidazole therapy. Additionally, the fact that P*nimB^G^* co-occurs with the Thr82Ile substitution in GyrA updates the paradigm for the global dissemination of epidemic strains^22^, indicating that the clinically apparent pandemic of fluoroquinolone-resistant *C. difficile* was co-mediated by resistance to metronidazole. These dual drug resistances likely afforded epidemic *C. difficile* an advantage in healthcare settings. Multiple lines of evidence support P*nimB^G^* as a cause of widespread metronidazole resistance, including our GWAS (Fig. 4a), a previously published GWAS of independent samples^45^, our documentation of the mutation emerging *in vivo* in a metronidazole-treated patient, and its mechanistic genetic validation as explained below.

Our analyses of population-level genomic data found *nimB* and *gyrA* variants are advantageous, as expected for drug resistance mutations. The two mutations were strongly linked, and phylogenetic analyses suggest that strains carrying both mutations spread more rapidly in healthcare settings (Fig. 4b-c). We found the combination of *nimB* and *gyrA* mutations in clade 2 and to a lesser extent clade 1 (Fig. 4b-d). This association could reflect niche specialization among the clades. Large scale surveys of *C. difficile* genomic data have identified high rates of carriage of both antimicrobial resistance determinants and toxin genes in clade 2^18, 45, 46^. This genetic cargo reflects adaptation of clade 2 to the antibiotic milieu of health care settings and pathogenicity in human diarrheal disease^18, 47^. Clade 5 is similarly associated with antibiotic resistance and increased virulence,^45^ but we did not identify the *nimB*-promoter mutation in our sample of clade 5 isolates (Fig. 4d). Clade 5 antimicrobial resistance determinants reflect the livestock and farming environments that are associated with this clade^18^ and absence of the *nimB*-promoter mutation from this clade could reflect the distinct antibiotic selection pressures encountered in these environments. It is also possible that the *nimB*-promoter mutation, and resulting heme-dependent metronidazole resistance, is a human-specific adaptation. Given the small number of clade 5 isolates included in our analyses, a larger sample size will be required to test this hypothesis. An alternative, but not mutually exclusive, explanation for the observed associations between the *nimB*-promoter mutation, the *gyrA* mutation and the clade 2 genetic background is epistasis, i.e., interactions among genetic loci that affect cell physiology. It is striking that the *nimB* mutation almost exclusively appears among strains with the *gyrA* mutation (Fig. 4b-d). A potential explanation for this phenomenon is that the *nimB*-promoter mutation and constitutive formation of *nimB* imposes a metabolic burden and fitness costs that might be ameliorated in strains with the *gyrA* mutation. However, 23468 and 23475, which are isogenic resistant and susceptible, did not show significant differences in growth rates (Extended Data Fig. 6). On the other hand, Thr82Ile mutation either marginally enhances fitness or has no fitness costs^48, 49^.

We discovered that *Cd*NimB acts as a nitroreductase both enzymatically and in cells and requires heme to effectively produce amino end products. According to the accepted model, Nim proteins convert nitroimidazoles to amine by rapidly transferring two electrons (a total of six) from unresolved cofactors to the nitro group^28, 38^. This requirement for heme may explain why Nim proteins were not reported to exhibit nitroreductase activity in conventional assays but was seen cellularly in heme-containing broths^28^. Nim proteins are structurally and functionally homologous to the flavoenzyme Anf3, in which neighboring heme and flavin binding domains facilitate rapid electron transfer between the two cofactors. Based on the Anf3 model (Fig. 3b), *Cd*NimB may either reduce nitroimidazole nitro-groups with two electrons from the individual heme molecules in the homodimeric protein, or the reduced hemes may transfer electrons to the flavin (e.g., FAD), which then transfers two electrons to the nitro-group. Further work will be required to test these hypotheses and determine the electron transfer path by which Nim proteins detoxify nitroimidazoles. Nonetheless, observations from this study contribute to resolving a long-standing debate on whether *nim* genes confer metronidazole resistance^37^. Indeed, *B. fragilis nimA* (OCL16839.1) conferred heme-dependent resistance when expressed in *C. difficile* (Extended Data Fig. 7a). Similarly, the *nimA* reference strain *B. fragilis* 638R/pIP417(*nimA*)^50^ showed heme-dependent resistance to metronidazole (MICs of 0.5-1 μg/ml and 4-8 μg/ml, without and with heme, respectively; Extended Data Fig. 7b). Thus, like *CdnimB*, other *nim* genes may be cryptic and expression of resistance may be influenced by their regulation, together with cellular amounts of heme and possibly flavin cofactors.

Overall, our study identified for the first time *nim* as a cryptic metronidazole resistance gene linked to the *C. difficile* epidemic. Noteworthy, intravenous metronidazole is still recommended with vancomycin for fulminant CDI by IDSA/SHEA and ESCMID^6, 7^ and metronidazole remains commonly used in the United States^51^ and in non-Western countries where it is still recommended to treat CDI^52, 53^. Therefore, this study provides several key insights that could be applied to modernize the use of metronidazole, as well as other CDI medications, through evidence-based approaches. To achieve this, rapid susceptibility testing, genomic surveillance, and mechanistic research will be required to delineate how resistance alleles influence clinical outcomes.

## Materials and Methods

### Bacterial strains

*C. difficile* strains used can be found in Supplementary Table 1. The geographic locations from which strains were isolated are specified: for GWAS strains were isolated in United States, Europe and Israel and other locations, as documented; other strains were from Texas Medical Center. Also included were reported from the United States Centers for Disease Control and Prevention Emerging Infections Program^54^. *B. fragilis* 638R/pIP417(strain DSM103646) and DSM2151 were from Leibniz Institute DSMZ; DSM2151 is an antibiotic susceptibility testing control strain. All strains of *C. difficile* or *B. fragilis* were cultured in pre-reduced Brain Heart Infusion (BHI) broth overnight at 37°C in a Whitley A35 anaerobic workstation (Don Whitley Scientific).

### Determination of minimum inhibitory concentrations (MICs) by agar dilution method

MICs were determined by the agar dilution method on BHI agar either with or without 5 µg/ml of porcine hemin (Alfa Aesar, catalog no. A11165) and doubling dilutions of metronidazole (0.125-32 µg/ml; from Acros, catalog no. 210340050), as previously described^20^. Metronidazole susceptibility tests in BHI were shown to be comparable to Wilkins Chalgren and Brucella agars^20^. Metronidazole and hemin stocks were prepared in dimethyl sulfoxide (DMSO) (Alfa Aesar, catalog no. 43998). Plates were incubated for up to 48 h and MICs recorded as the lowest concentration of drug that inhibited visible growth. MICs for 2-nitroimidazole (Sigma-Aldrich, catalog no.195650) were similarly performed.

### Generation of transposon mutagenesis library of metronidazole resistant *C. difficile*

Plasmid pRPF215, a *Himar1* mariner delivery vector^26^, was conjugated into R20291 from *E. coli* SD46 and R20291 transconjugants selected on BHI agars supplemented with 250 µg/ml cycloserine, 8 µg/ml of cefoxitin (Chem-Impex International, catalog no. 01490), and 15 µg/ml thiamphenicol (Sigma-Aldrich, catalog no. T0261). To generate Tn mutants, R20291-pRPF215 was grown overnight in BHI broth with 15 µg/ml thiamphenicol, then subcultured into BHI to OD600nm of 0.2. *Himar1* was induced with 100 ng/ml anhydrotetracycline (ATc) (Alfa Aesar, catalog no. J66688) for mutagenesis for overnight (15 h). Tn mutants were selected by spreading culture dilutions onto pre-reduced BHI agar containing 20 µg/ml lincomycin. After incubation (24 h), colonies were collected into 96 deep well plates, to amass 7,488 independent colonies. The library was screened for metronidazole-susceptible Tn mutants by growing overnight cultures in BHI containing lincomycin and spotting 1 μl inocula onto BHI agars containing 0.5 µg/ml metronidazole and 5 µg/ml of hemin using a 96-Well Bench Top Pipettor. Presumptive metronidazole-susceptible Tn mutants were confirmed by MIC testing and underwent genome and Sanger sequencing to confirm the insertion sites for *ermB*. Primers and strain constructs used are listed in the Supplementary Tables 2 and 3.

### Generation of knock out strain

To use CD2517.1 Type I toxin-antitoxin system as counter-selectable we modified vector pMTL-SC7215, essentially as previously described^55^, producing pMTL-SC7215-CD2517.1. To delete *nimB* by allelic exchange, a 1000 bp cassette of the upstream and downstream flanking regions were synthesized by Genscript, cloned into pMTL-SC7215-CD2517.1 and conjugated into R20291. After 8 serial passages in BHI, cultures were plated onto agars with ATc (70 ng/ml) and colonies were screened by PCR and Sanger sequencing for loss of *nimB*.

### Overexpression and knockdown of target genes

To overexpress *nimB* under its indigenous promoter, *nimB* together with its 504-bp upstream sequence was cloned between KpnI and BamHI sites of vector pRPF185, producing pRPF185-P*nimB*. Overexpression of *nimB* under anhydrotetracycline-inducible promoter (Ptet) was done by cloning the gene with its ribosome binding site into pRPF185, producing pRPF185-PATc*nimB*. Plasmid constructs were conjugated into *C. difficile* from *E. coli* SD46 and strains were grown in BHI broth with 15 µg/ml thiamphenicol. MICs were performed as previously indicated but in plates supplemented with thiamphenicol (15 µg/ml) and ATc (32 ng/ml) specifically for pRPF185-PATc*nimB*. Translational knockdown was performed as described^35^, whereby a 100-bp antisense RNA synthesized by Genscript, spanning 50-bp upstream and downstream of *nimB* from the start codon, was cloned into SphI and XhoI sites of vector pMSPT. Induction of gene knockdown was performed with 64 ng/ml of ATc. CRISPR interference was done as described^35^, whereby the guide RNA targeting *nimB* was synthesized by Genscript and cloned into the PmeI site of vector pXWxyl-dcas9. The guide RNA was induced with 1% xylose (Alfa Aesar, catalog no. A10643). Primers and strain constructs used are listed in the Supplementary Tables 2 and 3.

### Promoter strength analysis

The 504-bp upstream region of *nimB* was cloned into NheI and SacI sites of pDSW1728 vector to generate P*nimB*^G^::mCherryOpt or P*nimB*^T^::mCherryOpt from R20291 or CD196 respectively. Constructs and the empty vector control were conjugated into various *C. difficile* strains. Strains bearing the reporter construct and empty vector were grown to OD600nm ∼0.3, from overnight cultures. Each culture was either treated with DMSO control, metronidazole (2 µg/ml), metronidazole (2 µg/ml) plus hemin (5 µg/ml) or hemin alone (5 µg/ml) and incubated anaerobically for 1 h. Samples were then fixed as described previously^34^ and fluorescence measured in BioTek Synergy reader at excitation of 554 nm and emission of 610 nm. The fluorescence value for each sample was then normalized based on their culture densities (OD600nm values). Primers and strain constructs used are listed in Supplementary Tables 2 and 3.

### Construction and synthesis of Nim-like homologues

Genes encoding Anf3 and NimA from *A. vinelandii* and *B. fragilis*, respectively, were codon optimized for translation in *C. difficile* using JCat platform^56^. A synthetic ribosome-binding site for each gene was designed using RBS Calculator v2.1^57^ and placed ahead of the start codon and the ensemble synthesized by Genscript. They were then subcloned into pRPF185 vector and expressed from the *Ptet* promoter using 8 ng/ml of ATc. Primers and strain constructs used are listed in Supplementary Tables 2 and 3.

### RNA sequencing

R20291 was grown to OD600nm of 0.2 in BHI broth and exposed to DMSO control, metronidazole (2 µg/ml), metronidazole (2 µg/ml) plus hemin (5 µg/ml), or hemin alone (5 µg/ml) for 30 mins. RNA was extracted using Qiagen RNeasy miniKit (Qiagen, Valencia, CA) and treated with TURBO™ DNase (Thermofisher Scientific). RNA sequencing was then performed by GENEWIZ, LLC. (South Plainfield, NJ, USA), by 2×150 Paired End (PE) configuration by Illumina HiSeq. Raw RNA-seq data were uploaded onto galaxy platform (https://usegalaxy.org/). Quality control was carried out using FastQC (Babraham Bioinformatics) and multiFastQC. Trimming was done using Trim Galore to remove adapter sequences. Reads were mapped to R20291 reference FN545816.1 using the BWA-MEM program. Counts per Read was generated using SAM/BAM, while differential gene expression was done with Degust (http://degust.erc.monash.edu/) using edgeR (cut-offs abs log2 1 ≥ and FDR≤ 0.01). KEGG pathway analysis was done with ShinyGO platform^58^ with cut-offs log2 1≥ and FDR≤ 0.01.

### Reverse Transcription Quantitative Real-Time PCR

From total RNA, cDNA synthesis was done using qScript cDNA Supermix kit (Quantabio). qPCRs were carried out in 20 µL reactions, using qScript 1-Step SYBR Green qPCR kit (Quantabio) on a ViiA 7 Real-Time PCR System (Applied Biosystems); 16S rRNA was used for normalization. Gene expression was calculated by the standard ΔΔCT method. Constitutive gene expression analysis was calculated relative to metronidazole-susceptible strains. Primers used are listed in the Supplementary Table 2.

### Structural modeling of NimB

A homology model of R20291 *Cd*NimB was generated from *B. thetaiotaomicron* NimB dimeric protein structure with bound FAD (PDB 2FG9, resolution 2.20 angstrom). Prime software in the 2020-2 Schrödinger molecular modelling suite was used to build and refine the model^59, 60^. The *Bt*NimB and *Cd*NimB models were aligned to Anf3 nitrogenase from *A. vinelandii* (PDB 6RK0, resolution 0.99 angstrom^36^) using the Schrödinger software.

### Site directed mutagenesis of NimB

Heme-binding residues were predicted through alignment of *Cd*NimB with known heme-binding pyridoxamine 5’-phosphate oxidase family proteins, analysis in hemoquest and structural comparison with Anf3. Point mutations were generated in pRPF185-P*nimB* by standard DpnI mediated site-directed mutagenesis, variants were confirmed by Sanger sequencing and conjugated into *C. difficile*. Susceptibilities to metronidazole or 2-nitroimidazole were determined as described above. Primers used are listed in Supplementary Table 2.

### Purification of NimB proteins

Genes encoding NimB or NimB-His55Ala variant were cloned into pWL613a, a derivative of pET28b plasmid. They were transformed into *E. coli* BL21 (DE3), grown in Terrific Broth with 8µM hemin and induced with 0.4 mM IPTG at 14 ⁰C for 18 h. Harvested cultures were suspended in buffer (50 mM Tris-HCl (pH8.0), 0.5 M NaCl, 10 mM imidazole, 18% glycerol, 5mM *β-*mercaptoethanol, 1X protease inhibitor cocktail, 0.01 mg/ml DNase, 8 µM hemin and 5 mM MgCl2), lysed by French press 28,000 psi, His-tagged proteins were purified from Ni-NTA columns (Marvelgent Biosciences Inc), and dialyzed for 20 h at 4°C in buffer [20 mM Tris-HCl (pH8.0), 0.25 M NaCl, 18% glycerol and 5 mM *β-*mercaptoethanol].

### Analysis of heme binding by absorption spectroscopy

The ability of Nim to bind heme was determined using an established method^61^ in which the protein is titrated with heme. Nim proteins were purified as above, without hemin being added to the *E. coli* culture and lysis buffer. To determine heme binding, protein (10 µM) in buffer (20 mM Tris-HCl [pH 8.0], 0.25M NaCl, 18% w/v glycerol and 5 mM *β-*mercaptoethanol) was used to collect the absorbance spectra between 350-700 nm at 2 nm increments in polystyrene 24-well plates in the dark. The protein was titrated by stepwise addition of hemin from 1 mM stock in DMSO and absorbance spectra recorded after 5 min of incubation at room temperature using Synergy H1 BioTek microplate reader. Titration was continued until there was no further significant change in the spectra, indicating saturation. The spectra of the buffer containing hemin was recorded under the same conditions and subtracted from the protein samples. Heme binding was analyzed in plots of the change in absorbance at 416nm (Soret band) against the hemin concentration, and qualitatively by visualizing pooled samples.

### Nitroreductase enzymatic assay

NimB nitroreductase activity was assayed using substrates 4-nitrobenzoic acid, 2-nitroimidazole and metronidazole, with vancomycin as control; 4-nitrobenzoic acid and vancomycin were from Alfa Aesar, catalog no. A14738 and Sigma-Aldrich catalog no. V2002, respectively. The assay was done in an anaerobic chamber. Pyranose oxidase catalase was used to remove residual oxygen, as previously reported^36^. The reaction was performed in 50 mM KPO4 buffer with 10 µM hemin, 0.3 mM NADPH (Cayman, catalog no. 9000743), 10 µM FAD (Cayman, catalog no. 23386), 7.5 U/ml pyranose oxidase (Sigma-Aldrich, catalog no. P4234), 1 KU/ml catalase (Sigma-Aldrich, catalog no. C9322), 50 mM glucose and 5 mM substrate. The reaction mixture was pre-incubated at 37°C for 1 h before the addition of NimB, wildtype or His55Ala variant, at a final concentration of 9.7 µM and incubation continued for 2 h. Primary nitroaromatic amines were detected using the Bratton-Marshall assay reagents^62, 63^, while 2-aminoimidazole formation was further quantified by LC-MS/MS.

i. **Bratton-Marshall detection of nitroaromatic amines** In the anaerobic chamber, an equal volume of 20% cold Trichloroacetic acid (TCA) (Sigma-Aldrich, catalog no. T0699) was added to the reaction mixture, and after 30 mins on ice, 200 µL of supernatant was mixed with 25 µL of 0.1% sodium nitrite (Alfa Aesar, catalog no. A18668). After 10 mins, 25 µL of 0.5% w/v ammonium sulfamate (Alfa Aesar, catalog no. A17696) was added followed by 25 µL of 0.05% N-(1-Naphthyl)ethylenediamine dihydrochloride (Alfa Aesar, catalog no. A17164). After 20 mins, absorbance was measured at 540 nm with a VERSA Max microplate reader. Standard curves were generated using 4-aminobenzoic acid (Sigma-Aldrich, catalog no. A9878) and 2-aminoimidazole (Sigma-Aldrich, catalog no. CDS020502) were prepared in DMSO; no standard curve could be generated for 5-aminoimidazole as this chemical is unstable and commercially unavailable.
ii. **LC-MS/MS quantification of 2-aminoimidazole from enzymatic and cellular assays.** After incubation of the enzymatic or enzyme free reactions, samples were precipitated with cold TCA (10 % v/v). After 30 mins on ice, supernatants were collected, stored at -80°C and LC-MS/MS performed at Baylor College of Medicine NMR and Drug Metabolism Advanced Technology Core. This method was also used to measure formation of 2-aminoimidazole in cells as follows. Logarithmically growing cells (OD600nm ∼0.3) were concentrated 20-fold and exposed to DMSO control, 2-nitroimidazole (2 mM), 2-nitroimidazole (2 mM) plus hemin (5 µg/ml) or hemin alone (5 µg/ml) and incubated for 3 h anaerobically. After incubation, cells were treated with TCA (10 % (v/v) and supernatants recovered. For LC-MS/MS, samples were diluted in methanol. 2-aminoimidazole was resolved, identified, and quantified by UHPLC coupled with Q Exactive Orbitrap MS (Thermo Fisher Scientific) equipped with 50 mm x 4.6 mm column (XDB C-18, Agilent Technologies). MS data were acquired from 50 to 1000 Da in profile mode and reference ions at *m/z* 371.1012. A standard curve was generated for 2-aminoimidazole to determine concentrations in samples and results were normalized per protein content.

### Reference guided assembly

We used reference-guided assemblies instead of *de novo* assemblies to better query variations in intergenic regions. We conducted reference-guided assembly on 365 U.S. *C. difficile* isolates and 136 global, clade 2, 027/BI/NAP1 *C. difficile* isolates from He et al.^22^ Raw sequencing data were assembled using a reference-guided assembly pipeline (https://github.com/pepperell-lab/RGAPepPipe.git). Quality of the sequencing data was assessed using FastQC v0.8.11^64^, reads were trimmed using TrimGalore v0.6.4 (https://github.com/FelixKrueger/TrimGalore) and then aligned to R20291 reference sequence FN545816.1, using BWA MEM v0.7.12^65^. Alignments were processed using samtools v1.3.1^66^, and Picard v1.183 (http://broadinstitute.github.io/picard/) was used to remove duplicates and add read information. Reads were realigned using GATK v3.5^67^ and variants were identified using Pilon v1.16^68^. Finally, assembly quality was assessed using Qualimap BamQC^69^. We discarded 18 isolates with poor quality assemblies, evaluated by the percent of reads aligned to the reference as < 70%. The final dataset comprised of 348 *C. difficile* isolates (347 isolates + 1 reference); average read depth values of the dataset used in our analyses ranged from 31X to 149X; median coverage ranged from 100X to 300X; and all samples analyzed had >70% alignment to the reference.

### Phylogeny

A phylogenetic tree of the whole-genome sequences of our isolates was inferred using RAxML v8.2.3^70^ using the maximum-likelihood method under the General Time Reversible (GRT) model of nucleotide substitution and the CAT approximation of rate heterogeneity. The tree was visualized in R using ggtree^71^. Clade typing was verified using clade representatives from Dingle et al.^72^ A phylogenetic tree made by combining clade 2 isolates from this study and the whole-genome sequences assembled from He et al.^22^ was also inferred using RAxML and the GTRCAT model.

### Fst outlier analysis

To identify genetic variants associated with our phenotype of interest, we performed an Fst outlier analysis between susceptible and heme-dependent resistant populations. From the whole-genome alignment of all our isolates we created a VCF using SnpSites v2.4.1^73^ and calculated Weir and Cockerham’s Fst (wcFst) for bi-allelic SNPs using vcflib (https://github.com/vcflib/vcflib). To identify wcFst outliers we repeated our analysis 100X using randomly assigned phenotypes and used the maximum wcFst value observed in this null distribution as a significance cut-off.

### Homoplasy analysis

Homoplasy analysis was conducted using Treetime^74^ to investigate *gyrA* and *nimB* SNPs for signs of selection.

## Acknowledgements

This work was funded by R01AI139261 from the National Institute of Allergy and Infectious Diseases at the National Institutss of Health. The funders had no role in study design, data collection, interpretation of the findings, or in the writing and submission of the manuscript. We are grateful for the receipt of clinical strains from Amos Adler, Tel Aviv University, Israel; Paola Mastrantonio, Istituto Superiore di Sanità, Italy; Dale Gerding, Loyola University Medical Center, United States; Scott Curry, University of Pittsburgh School of Medicine, United States; Mary-Beth Dorr, MODIFY trials program, of Merck & Co., Inc; various participants of the Pan-European Longitudinal Surveillance of Antibiotic Resistance among Prevalent *Clostridium difficile* Ribotypes’ Study Group; BEI Resources, National Institute of Allergy and Infectious Diseases at the National Institutes of Health; and United States Centers for Disease Control and Prevention Emerging Infections Program. We are also grateful to Jordan May for technical assistance with genome extractions.

## Author Contributions

J.G.H initially conceived the project. A.O.O., C.D., M.A.Y., A.S., M.A.T., C.S.P., and J.G.H conceived, conducted experimental design, execution, and data analysis, and wrote the original draft. All authors reviewed the manuscript. W.S., A.D., and K.L.P. established genome extraction protocols; W.S. and A.D. contributed to establishing susceptibility testing protocols. K.E.H. conducted molecular modeling of NimB. K.W.G and A.G.L. provided clinical isolates from Houston and reviewed susceptibility testing results. M.H.W. and J.F. provided clinical isolates from Europe.

## EXTENDED FIGURES

**Extended Figure 1.**
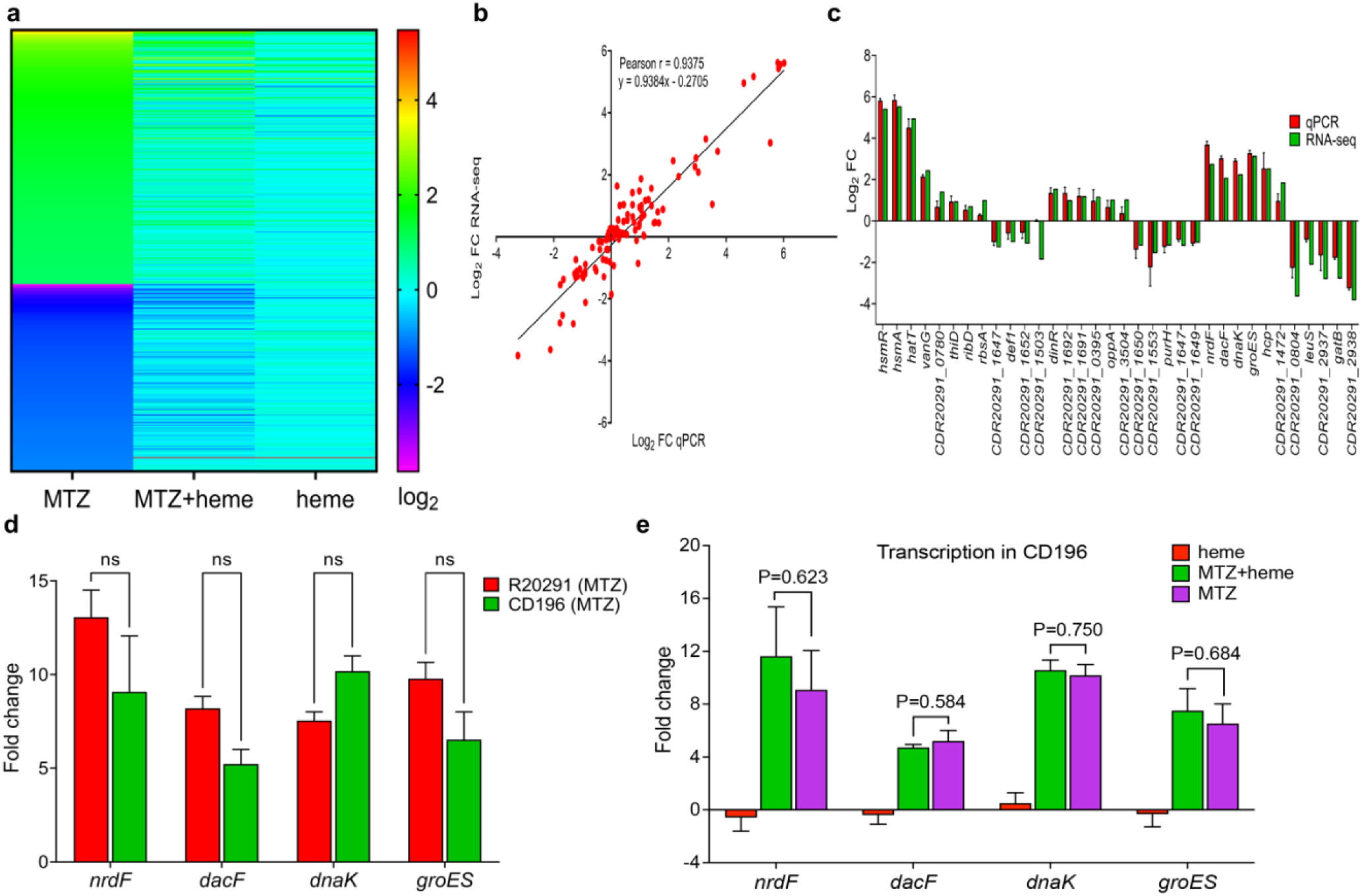
The effect of heme on the transcriptome of epidemic R20291 in response to metronidazole (MTZ) toxicity, and comparison to non-epidemic CD196. **(a)** Heat map showing pattern of differentially expressed genes (DEG) in R20291 in MTZ (2 μg/ml), MTZ plus heme (5 μg/ml) or heme alone. DEGs were determined using an FDR cutoff of ≤ 0.01 and an absolute value of Log2 Fold Change *≥* 2, relative to DMSO exposed culture. (b) Pearson correlation plot from a panel of 34 genes (20 upregulated and 11 downregulated). The correlation coefficient (R^2^) was 0.94 (P<0.0001). Of the three biological replicates from RNAseq, one was discarded due to poor quality in FASTQC, but gene expression was validated by qPCR using three biological replicates. (c) Transcription of selected genes from the above RNAseq and qPCR data. (d) In the absence of heme, CD196 and R20291 display a similar transcription pattern. (e) In MTZ-susceptible CD196, heme does not attenuate MTZ responsive gene transcription, unlike in R20291 (see Fig. 1d of the main text). Statistical significance was assessed by unpaired t-test, ns= not significant; data in d and e, represent the means ± standard error of mean from three biological replicates.

**Extended Figure 2.**
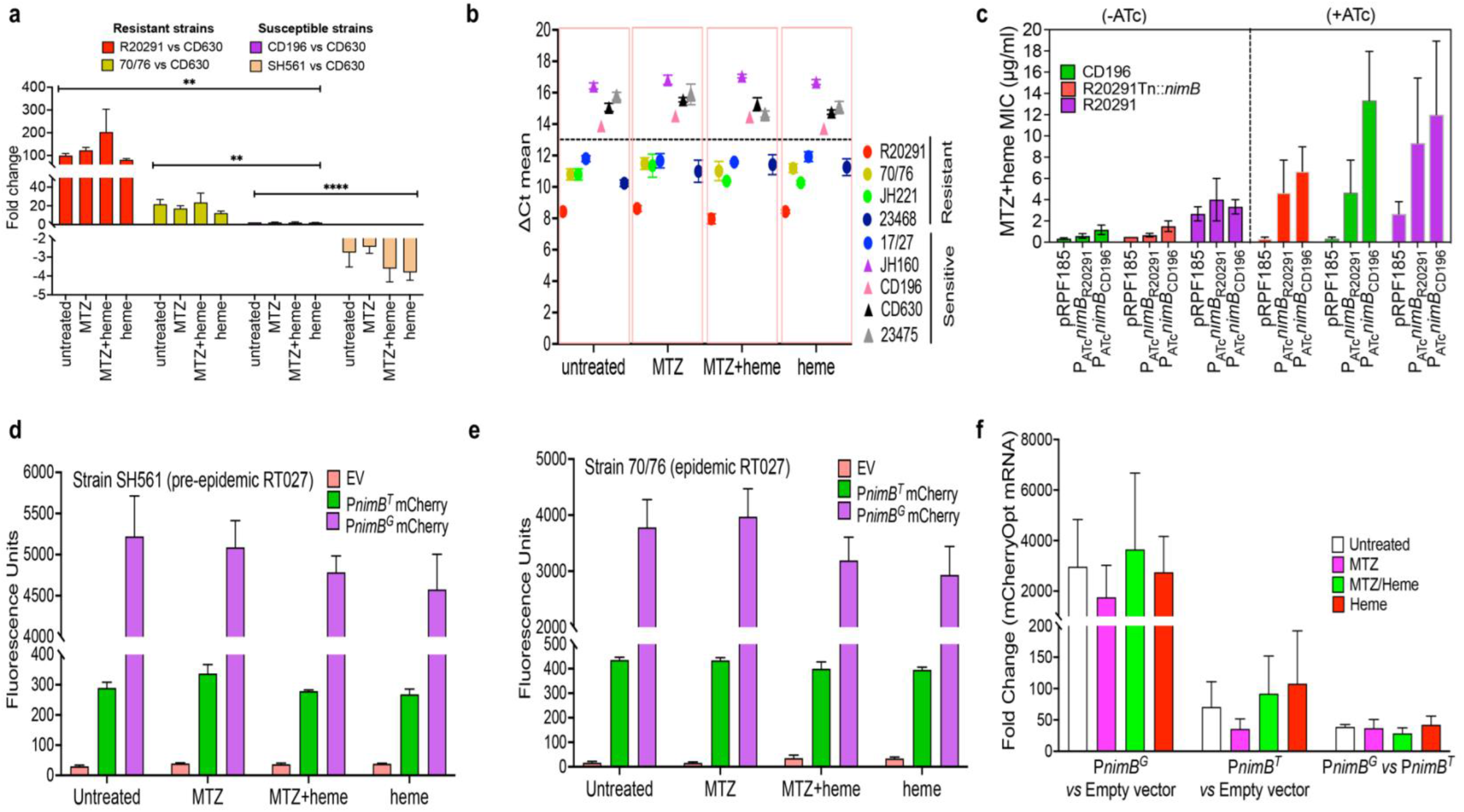
Identification and validation of *nimB* as a mechanism of heme-dependent resistance to metronidazole (MTZ). **(a)** qPCR analysis of *nimB* transcription in resistant and susceptible strains with respect to the susceptible lab strain CD630, under different conditions. Data represent the means ± standard error of mean of three biological replicates; statistical significance was assessed for all compiled data on each strain by Welch and Brown-Forsythe ANOVA with Dunnett’s Multiple Comparison in GraphPad Prism 9.4.1; p<0.0021, **; p<0.0001, ****. **(b)** Relative cycle threshold (Ct) values of *nimB* transcripts among resistant (R20291, 70/76, 23468) and susceptible (CD196, CD630, SH561, 23475) isolates under different conditions. Ct values were normalized to the housekeeping 16S rRNA. Data represent the mean ± standard error of mean from three biological replicates. **(c)** MICs of MTZ upon overexpression of CD196 and R20291 *nimB* ORFs under the control of the ATc-inducible promoter (P*tet*). Expression was induced by 64 ng/ml ATc. **(d, e)** Further comparison of promoter strengths of P*nimB^G^* and P*nimB*^T^ based on mCherryOpt transcription in SH561 and 70/76, susceptible and resistant, respectively. Fluorescence was normalized to OD600nm culture density; MTZ MICs in heme were 0.25 μg/ml and 8 μg/ml against SH561 and 70/76, respectively. The fluorescence of P*nimB^G^* was higher across all test conditions, indicating constitutive expression, supporting findings in Fig. 2f of the main text. **(f)** mCherry mRNA levels were also higher with P*nimB^G^* than *Pnim^T^*; expressions were performed in CD196.

**Extended Figure 3.**
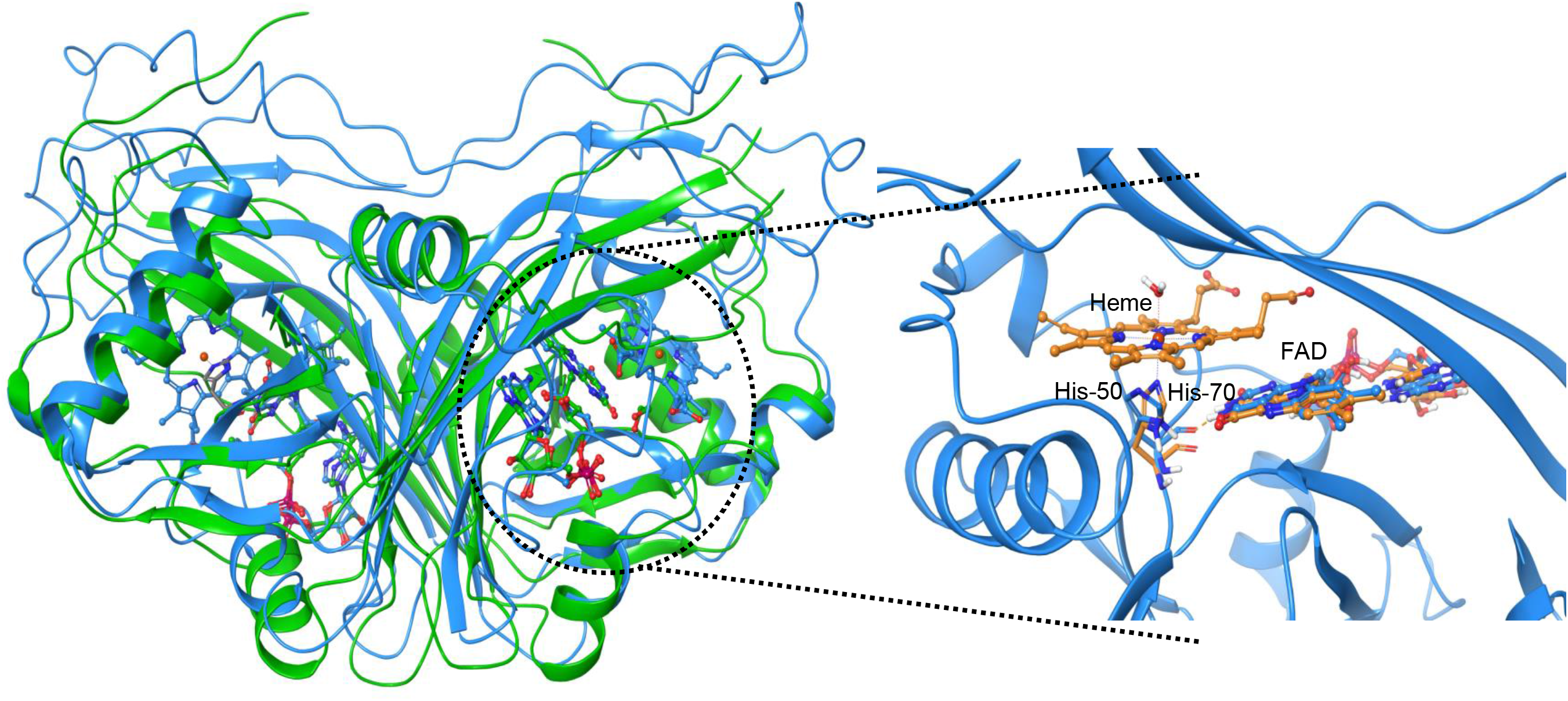
Overlay of *Bacteroides thetaiotaomicron* NimB crystal structure (PDB 2FG9, blue ribbons, resolution 2.20 angstrom) with Anf3 (PDB 6RK0, green ribbons, resolution 0.99 angstrom). Both previously published proteins are dimers and closely align; the close-up shows histidine-50 in *Bt*NimB is homologous to the heme binding proximal ligand histidine-70 of Anf3. *Bt*NimB was co-crystalized with FAD, while Anf3 was co-crystalized with both heme and FAD; the FAD cofactors of both proteins occupy the same relative positions in their cognate proteins. The *Bt*NimB structure was used to generate a homology model of *Cd*NimB in Figure 3b of the main text.

**Extended Figure 4.**
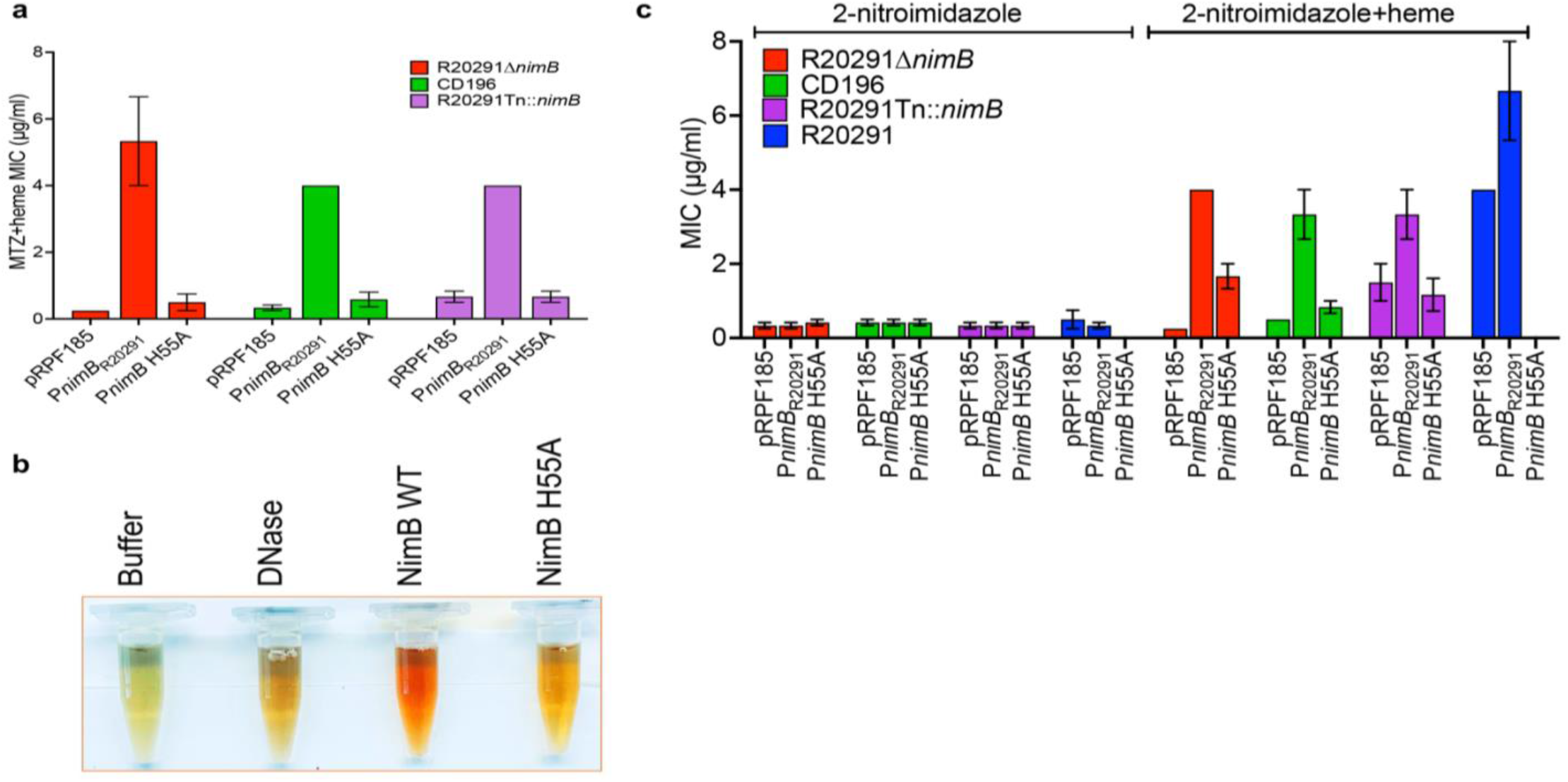
The effect of heme binding by *Cd*NimB on heme-dependent metronidazole (MTZ) resistance was examined. **(a)** The His55Ala (i.e., H55A) mutation does not confer resistance when expressed in various MTZ-susceptible strains; wildtype or H55A variant was expressed in plasmid pRPF185 under P*nimB^G^* promoter in the indicated strains (MTZ-susceptible CD196, R20291-*Tn::nimB* and R20291Δ*nimB*) and resistant R20291. MICs are shown as means ± standard error of mean from three biological replicates. **(b)** Qualitative evaluation of heme binding by *Cd*NimB and its H55A variant. Wildtype *Cd*NimB-heme forms a characteristic reddish-brown color of heme-protein complexation, but the H55A variant exhibits a more yellow color with heme, similar to the non-heme-binding DNase control. **(c)** Analysis of MICs of 2-nitroimidazole in the absence and presence of heme against abovementioned MTZ-susceptible and resistant strains.

**Extended Figure 5.**
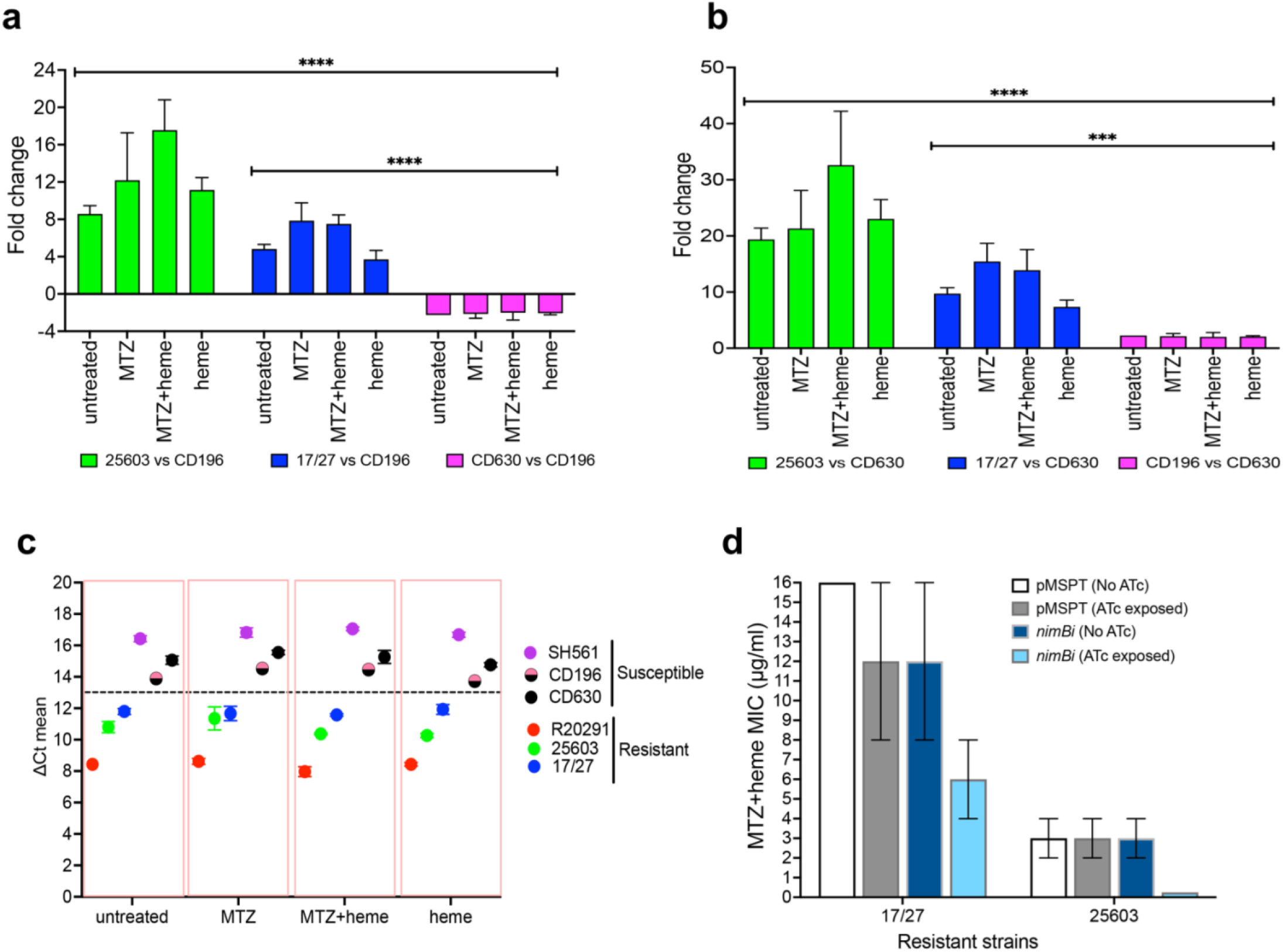
Transcriptional analysis of *nimB* in two heme-dependent metronidazole-resistant strains. **(a, b)** qPCR analysis of *nimB* transcription in strains 17/27 and 25603 with respect to the susceptible strains CD196 or the lab strain CD630, under different conditions. Data represent the means ± standard error of mean of three biological replicates; statistical significance was assessed for all compiled data on each strain by Welch and Brown-Forsythe ANOVA with Dunnett’s Multiple Comparison in GraphPad Prism 9.4.1; p<0.0002, ***; p<0.0001, ****. **(c)** Relative cycle threshold (Ct) values of *nimB* transcripts in among resistant (17/27 and 25603, with control R20291), compared to susceptible (CD196, CD630, SH561) isolates under different conditions. Ct values were normalized to the housekeeping 16S rRNA. Data represent the mean ± standard error of mean from three biological replicates. **(d)** Genetic silencing of *nimB* reduced MTZ resistance in 17/27 or completely reverses resistance in 25603. *nimB* was silenced by an antisense RNA (asRNA), which was induced by anhydrotetracycline (64 ng/ml) from P*tet* promoter in vector pMSPT.

**Extended Figure 6.**
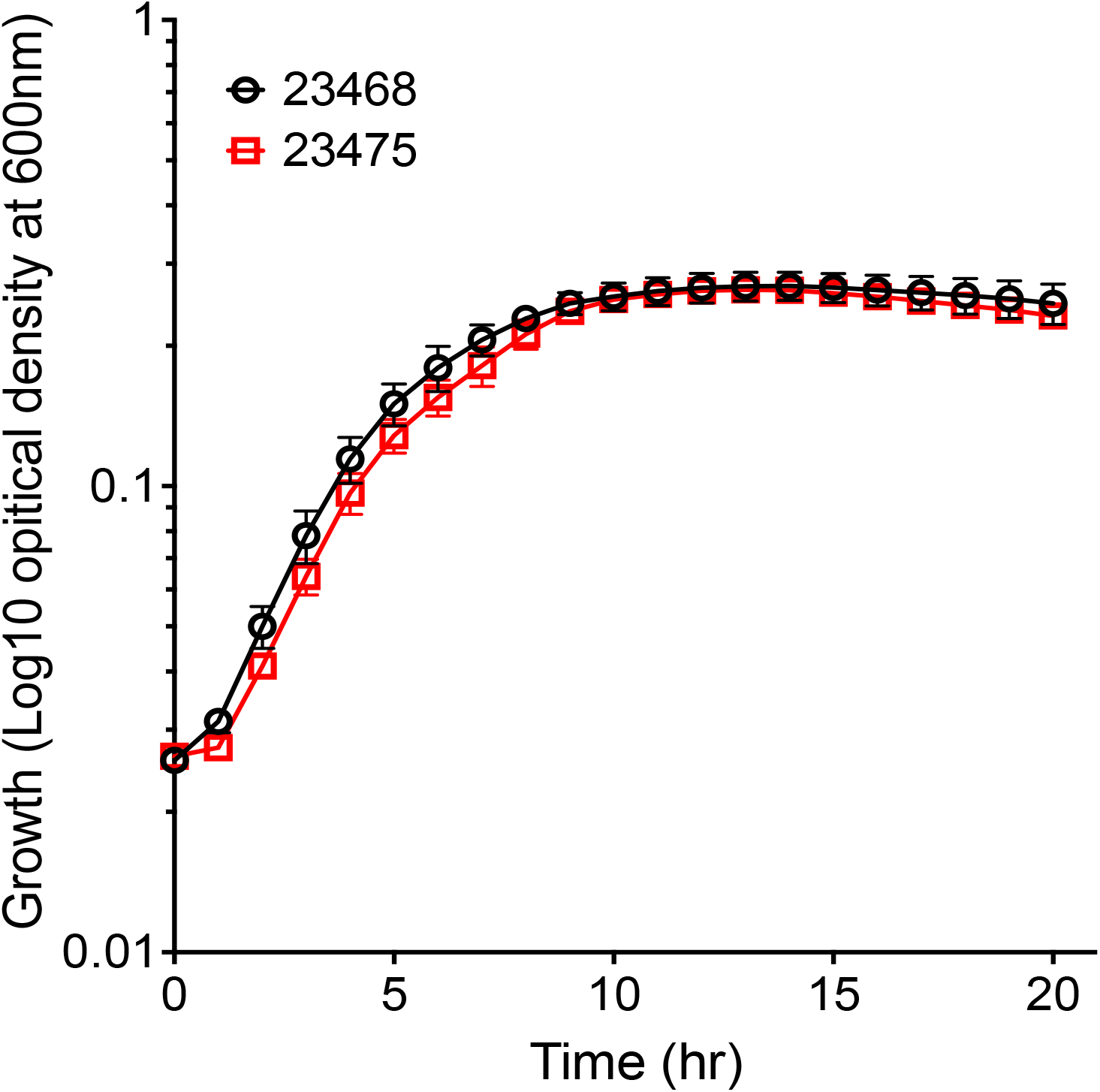
Comparison of growth of isogenic strains. 23475 (susceptible) and 23468 (resistant) show comparable growth curves in BHI broths; shown are three biological replicates with standard deviation error bars.

**Extended Figure 7.**
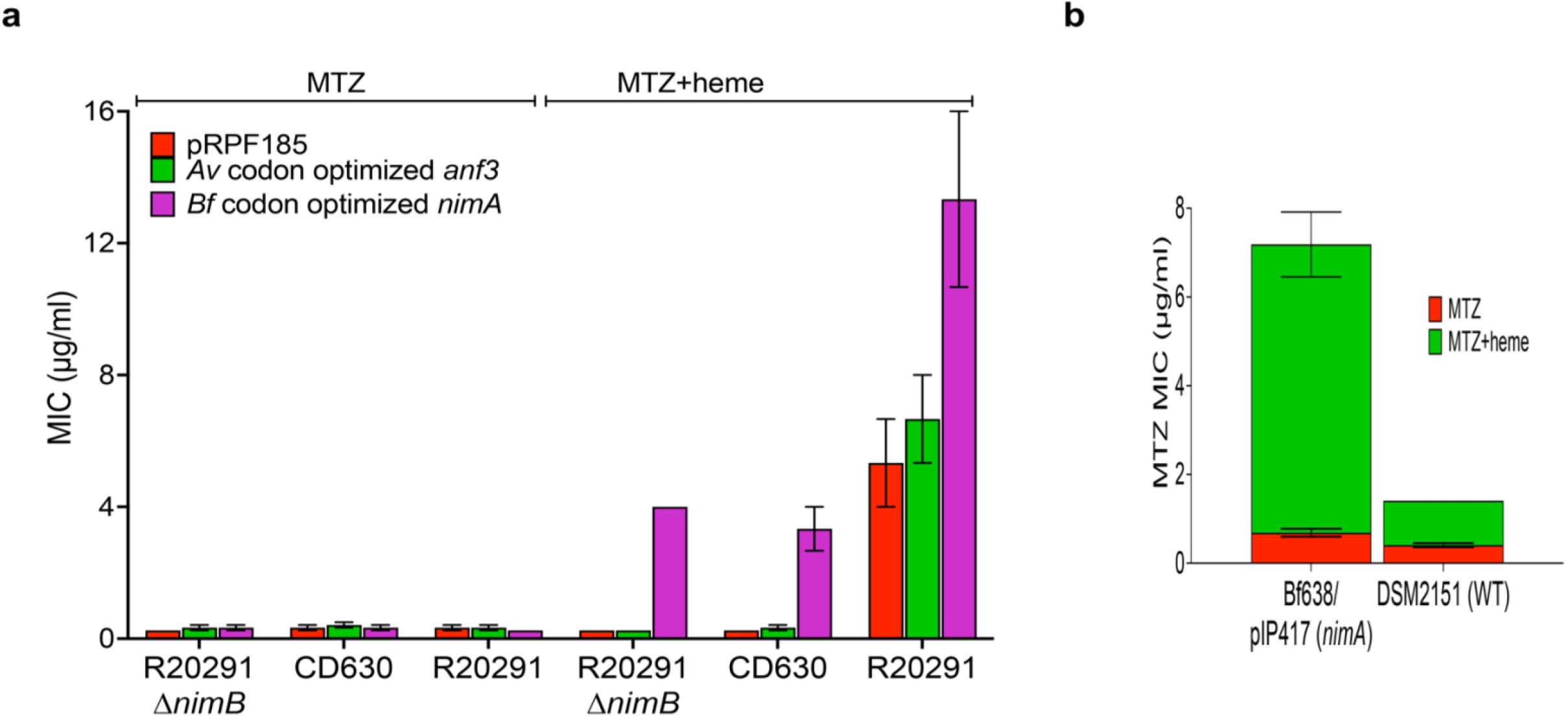
Heme-dependent metronidazole (MTZ) resistance is exhibited by *Bacteroides fragilis* NimA. **(a)** Recombinant NimA from *B. fragilis* (*Bf*) confers heme-dependent metronidazole (MTZ) resistance in susceptible **strains and increases resistance in R20291**. While Nim proteins (e.g., *Bf*NimA and *C. difficile* NimB are structurally related to Anf3, the latter did not confer heme-dependent MTZ resistance). Genes encoding *Bf* and *Av* proteins were codon-optimized for expression in *C. difficile*. **(b)** Susceptibility testing of *B. fragilis* known metronidazole-resistant strain Bf638/ pIP417 (*nimA*) and susceptible in the presence and absence of heme. Bf638/ pIP417 (*nimA*) showed that its resistance to metronidazole is heme-dependent; Bf638/ pIP417 (*nimA*) encodes a plasmid-borne *nimA*. DSM2151is a *B. fragilis* susceptibility testing control strain that is susceptible to metronidazole.

