## Supplementary Tables 1 - 3 for "Decoding a cryptic mechanism of metronidazole resistance among globally disseminated fluoroquinolone-resistant *Clostridioides difficile*"

**Supplementary Table 1.** Strains of *C. difficile* studied; MICs are graphed in Figure 1a of the main text.

| Isolate | Strain | Ribotype | Phenotype | MTZ MIC (mg/L) without heme | MTZ MIC (mg/L) with heme | Country of isolation (where known) | Isolation date (where known) | gyrA (mutant) | nimB (PnimBG) | SNP_Presence | Clade | Mutation analysis | Accession no., Bioproject ID |
|---|---|---|---|---|---|---|---|---|---|---|---|---|---|
| JH176 | SH758 | NA | S | 0.5 | 0.5 | USA | 5/3/17 | FALSE | FALSE | none | 2 | FST, GWAS | This study |
| JH177 | MT5054 | NA | S | 0.25 | 0.25 | USA | 5/3/17 | FALSE | FALSE | none | 1 | FST, GWAS | This study |
| JH179 | SH797 | NA | HMR | 0.5 | 8 | USA | 6/6/17 | TRUE | TRUE | gyrA and nimB | 2 | FST, GWAS | This study |
| JH206 | MT5094 | NA | S | 0.5 | 0.25 | USA | 7/8/17 | FALSE | FALSE | none | 1 | FST, GWAS | This study |
| JH275 | SH801 | NA | S | 0.25 | 0.25 | USA | 6/7/17 | FALSE | FALSE | none | 1 | FST, GWAS | This study |
| JH205 | SH1138 | FP435 | S | 0.5 | 0.5 | USA | 1/19/18 | FALSE | FALSE | none | 1 | FST, GWAS | This study |
| JH204 | SH1158 | FP423 | S | 0.5 | 1 | USA | 2/4/18 | FALSE | FALSE | none | 2 | FST, GWAS | This study |
| JH175 | MT5063 | FP310 | HMR | 0.25 | 4 | USA | 5/8/17 | TRUE | FALSE | gyrA only | 1 | FST, GWAS | This study |
| JH268 | MT3236 | 078-134 | S | 0.25 | 0.25 | USA | 8/17/11 | FALSE | FALSE | none | 5 | FST, GWAS | This study |
| JH267 | MT2694 | 078-132 | S | 0.5 | 0.25 | USA | 1/30/15 | FALSE | FALSE | none | 5 | FST, GWAS | This study |
| JH266 | MT3763 | 078-129 | S | 0.25 | 0.25 | USA | 9/19/15 | FALSE | FALSE | none | 5 | FST, GWAS | This study |
| JH265 | MT3245 | 078-127 | S | 0.25 | 0.25 | USA | 8/17/11 | FALSE | FALSE | none | 5 | FST, GWAS | This study |
| JH264 | MT2748 | 078-126 | S | 0.25 | 0.25 | USA | 11/30/14 | FALSE | FALSE | none | 5 | FST, GWAS | This study |
| JH276 | MT263 | 078-126 | S | 0.5 | 0.5 | USA | 9/30/11 | FALSE | FALSE | none | 5 | FST, GWAS | This study |
| JH277 | MT2717 | 078-126 | S | 0.25 | 0.5 | USA | 11/30/14 | FALSE | FALSE | none | 5 | FST, GWAS | This study |
| JH278 | MT2022 | 078-126 | S | 0.25 | 0.25 | USA | 6/22/13 | FALSE | FALSE | none | 5 | FST, GWAS | This study |
| JH280 | MT2127 | 078-126 | S | 0.25 | 0.25 | USA | 8/20/13 | NA | NA | NA | NA | FST, GWAS | This study |
| JH347 | MT4899 | 078-126 | S | 0.5 | 0.25 | USA | 2011 | FALSE | FALSE | none | 5 | FST, GWAS | This study |
| BAA-1875 | BAA-1875 | 078 | S | 0.5 | 0.25 | NA | NA | NA | NA | NA | NA | NA | NA |
| AR-1077 | AR-1077 | 078 | S | 0.25 | 0.25 | USA | 2016 | FALSE | FALSE | none | NA | Published genome | PRJNA577141 |
| AR-1083 | AR-1083 | 078 | S | 0.25 | 0.25 | USA | 2016 | FALSE | FALSE | none | NA | Published genome | PRJNA577141 |
| JH282 | 23381 | 078 | HMR | 0.25 | 2 | NA | NA | FALSE | FALSE | none | 5 | FST, GWAS | This study |
| JH285 | 27439 | 078 | S | 0.25 | 0.25 | NA | NA | FALSE | FALSE | none | 5 | FST, GWAS | This study |
| JH286 | 25944 | 078 | S | 0.25 | 0.25 | NA | NA | FALSE | FALSE | none | 5 | FST, GWAS | This study |
| JH330 | NR-49310 | 078 | S | 0.5 | 0.5 | USA | 1/27/17 | FALSE | FALSE | none | 5 | FST, GWAS | This study |
| JH434 | L,13.7933036 | 078 | S | 0.25 | 0.25 | Greece | 2012 | FALSE | FALSE | none | 5 | FST, GWAS | This study |
| JH438 | L,13.7933058 | 078 | S | 0.25 | 0.25 | Austria | 2011-2012 | FALSE | FALSE | none | 5 | FST, GWAS | This study |
| JH440 | L,13.7933149 | 078 | S | 0.25 | 0.25 | Spain | 2012 | FALSE | FALSE | none | 5 | FST, GWAS | This study |
| JH472 | L,13.7933425 | 078 | S | 0.25 | 0.25 | Italy | 2011-2012 | FALSE | FALSE | none | 5 | FST, GWAS | This study |
| JH479 | L,13.7933461 | 078 | S | 0.25 | 0.25 | UK | 2012-2013 | FALSE | FALSE | none | 5 | FST, GWAS | This study |
| JH489 | L,13.7933876 | 078 | S | 0.25 | 0.25 | UK | 2013 | FALSE | FALSE | none | 5 | FST, GWAS | This study |
| JH510 | L,12.7903869 | 078 | S | 0.25 | 0.25 | Hungary | 2014 | FALSE | FALSE | none | 5 | FST, GWAS | This study |
| JH545 | L,16.7569759 | 078 | S | 0.25 | 0.25 | Portugal | NA | FALSE | FALSE | none | 5 | FST, GWAS | This study |
| JH546 | L,16.7569864 | 078 | S | 0.25 | 0.25 | Switzerland | NA | FALSE | FALSE | none | 5 | FST, GWAS | This study |
| JH547 | L,16.7569846 | 078 | S | 0.25 | 0.25 | Netherlands | NA | FALSE | FALSE | none | 1 | FST, GWAS | This study |
| AR-1068 | AR-1068 | 056 | HMR | 0.25 | 1 | USA | 2016 | FALSE | FALSE | none | NA | Published genome | PRJNA577141 |
| AR-1079 | AR-1079 | 056 | S | 0.25 | 0.25 | USA | 2016 | FALSE | FALSE | none | NA | Published genome | PRJNA577141 |
| AR-1082 | AR-1082 | 054 | S | 0.25 | 0.25 | USA | 2016 | FALSE | FALSE | none | NA | Published genome | PRJNA577141 |
| AR-1088 | AR-1088 | 054 | S | 0.25 | 0.25 | USA | 2016 | FALSE | FALSE | none | NA | Published genome | PRJNA577141 |
| JH328 | NR-49325 | 054 | S | 0.25 | 0.25 | USA | 2011 | FALSE | FALSE | none | 1 | FST, GWAS | This study |
| JH329 | NR-49327 | 054 | S | 0.5 | 0.5 | USA | 2011 | FALSE | FALSE | none | 1 | FST, GWAS | This study |
| JH478 | L,13.7933418 | 054 | S | 0.25 | 0.25 | Italy | 2012 | FALSE | FALSE | none | 1 | FST, GWAS | This study |
| JH173 | MT5056 | 053-163 | HMR | 0.25 | 4 | USA | 5/7/17 | TRUE | FALSE | gyrA only | 1 | FST, GWAS | This study |
| JH327 | NR-49314 | 047 | S | 0.5 | 0.25 | USA | 2011 | FALSE | FALSE | none | 4 | FST, GWAS | This study |

| | | | | | | | | | | | | | |
|---|---|---|---|---|---|---|---|---|---|---|---|---|---|
| JH262 | MT1411 | 039 | HMR | 0.25 | 2 | USA | 11/30/12 | TRUE | TRUE | gyrA and nimB | 2 | FST, GWAS | This study |
| JH261 | MT201 | 036 | HMR | 0.25 | 2 | USA | 9/27/11 | TRUE | TRUE | gyrA and nimB | 2 | FST, GWAS | This study |
| JH260 | MT207 | 033 | HMR | 0.5 | 2 | USA | 9/30/11 | TRUE | TRUE | gyrA and nimB | 2 | FST, GWAS | This study |
| R20291 | R20291 | 027 | HMR | 0.25 | 2 | UK | 2006 | TRUE | TRUE | gyrA and nimB | 2 | FST, GWAS | FN545816.1 |
| BAA-1803 | BAA-1803 | 027 | HMR | 0.5 | 2 | NA | NA | NA | NA | NA | NA | NA | NA |
| CD196 | CD196 | 027 | S | 0.25 | 0.25 | France | 1985 | FALSE | FALSE | none | none | Genome analysis | PRJNA224116 |
| AR-1067 | AR-1067 | 027 | HMR | 0.25 | 2 | USA | 2016 | TRUE | TRUE | gyrA and nimB | NA | Published genome | PRJNA577141 |
| AR-1071 | AR-1071 | 027 | HMR | 0.25 | 2 | USA | 2016 | TRUE | TRUE | gyrA and nimB | NA | Published genome | PRJNA577141 |
| AR-1072 | AR-1072 | 027 | HMR | 0.25 | 2 | USA | 2016 | TRUE | TRUE | gyrA and nimB | NA | Published genome | PRJNA577141 |
| AR-1076 | AR-1076 | 027 | HMR | 0.25 | 2 | USA | 2016 | TRUE | TRUE | gyrA and nimB | NA | Published genome | PRJNA577141 |
| AR-1092 | AR-1092 | 027 | HMR | 0.25 | 2 | USA | 2016 | TRUE | TRUE | gyrA and nimB | NA | Published genome | PRJNA577141 |
| AR-1095 | AR-1095 | 027 | HMR | 0.25 | 2 | USA | 2016 | TRUE | TRUE | gyrA and nimB | NA | Published genome | PRJNA577141 |
| 70/75 | 70/75 | 027 | HMR | 1 | 8 | NA | NA | NA | TRUE | NA | NA | NA | This study |
| 70/76 | 70/76 | 027 | HMR | 1 | 8 | NA | NA | TRUE | TRUE | gyrA and nimB | NA | Genome analysis | This study |
| JH142 | MT3723 | 027 | HMR | 1 | 8 | USA | NA | TRUE | TRUE | gyrA and nimB | 2 | FST, GWAS | This study |
| JH148 | MT3678 | 027 | HMR | 0.5 | 2 | USA | 8/21/15 | TRUE | TRUE | gyrA and nimB | 2 | FST, GWAS | This study |
| JH149 | MT240 | 027 | HMR | 0.25 | 16 | USA | 9/30/11 | TRUE | TRUE | gyrA and nimB | 2 | FST, GWAS | This study |
| JH150 | SH1120 | 027 | HMR | 1 | 2 | USA | 1/13/18 | TRUE | TRUE | gyrA and nimB | 2 | FST, GWAS | This study |
| JH151 | MT4968 | 027 | HMR | 0.5 | 4 | USA | 3/3/17 | TRUE | TRUE | gyrA and nimB | 2 | FST, GWAS | This study |
| JH152 | SH691 | 027 | HMR | 0.5 | 4 | USA | 3/16/17 | TRUE | TRUE | gyrA and nimB | 2 | FST, GWAS | This study |
| JH153 | MT4883 | 027 | HMR | 0.25 | 2 | USA | 1/13/17 | TRUE | TRUE | gyrA and nimB | 2 | FST, GWAS | This study |
| JH154 | MT4887 | 027 | HMR | 1 | 16 | USA | 1/14/17 | TRUE | TRUE | gyrA and nimB | 2 | FST, GWAS | This study |
| JH155 | SH552 | 027 | HMR | 0.5 | 16 | USA | 12/30/16 | TRUE | TRUE | gyrA and nimB | 2 | FST, GWAS | This study |
| JH156 | MT4828 | 027 | HMR | 1 | 4 | USA | 12/25/16 | TRUE | TRUE | gyrA and nimB | 2 | FST, GWAS | This study |
| JH157 | SH648 | 027 | HMR | 0.25 | 2 | USA | 2/25/17 | TRUE | TRUE | gyrA and nimB | 2 | FST, GWAS | This study |
| JH158 | MT4803 | 027 | HMR | 0.5 | 16 | USA | 12/14/16 | TRUE | TRUE | gyrA and nimB | 2 | FST, GWAS | This study |
| JH159 | SH515 | 027 | HMR | 0.25 | 4 | USA | 12/9/16 | TRUE | TRUE | gyrA and nimB | 2 | FST, GWAS | This study |
| JH160 | SH561 | 027 | S | 0.5 | 0.25 | USA | 1/11/17 | FALSE | FALSE | none | 2 | FST, GWAS | This study |
| JH181 | MT3135 | 027 | HMR | 0.5 | 2 | USA | 4/25/14 | TRUE | TRUE | gyrA and nimB | 2 | FST, GWAS | This study |
| JH182 | MT2042 | 027 | S | 0.25 | 0.5 | USA | 7/11/13 | FALSE | FALSE | none | 1 | FST, GWAS | This study |
| JH183 | MT2202 | 027 | S | 0.25 | 0.25 | USA | 8/12/13 | FALSE | FALSE | none | 2 | FST, GWAS | This study |
| JH184 | MT211 | 027 | S | 0.25 | 0.25 | USA | 9/30/11 | FALSE | FALSE | none | 1 | FST, GWAS | This study |
| JH185 | MT271 | 027 | S | 1 | 1 | USA | 9/30/11 | FALSE | FALSE | none | 2 | FST, GWAS | This study |
| JH186 | MT2756 | 027 | S | 1 | 1 | USA | 11/30/14 | FALSE | FALSE | none | 1 | FST, GWAS | This study |
| JH187 | MT749 | 027 | S | 1 | 1 | USA | 3/22/12 | FALSE | FALSE | none | 1 | FST, GWAS | This study |
| JH188 | MT322 | 027 | S | 1 | 1 | USA | 9/30/11 | FALSE | FALSE | none | 1 | FST, GWAS | This study |
| JH189 | MT3139 | 027 | S | 1 | 1 | USA | 10/14/14 | FALSE | FALSE | none | 1 | FST, GWAS | This study |
| JH207 | .difficile 1675- | 027 | S | 0.25 | 0.25 | NA | Feb-88 | FALSE | FALSE | none | 2 | FST, GWAS | This study |
| JH208 | .difficile 4173- | 027 | S | 0.25 | 0.5 | NA | Oct-90 | FALSE | FALSE | none | 2 | FST, GWAS | This study |
| JH209 | .difficile 4196- | 027 | S | 0.25 | 0.25 | NA | Nov-90 | FALSE | FALSE | none | 2 | FST, GWAS | This study |
| JH210 | CD43 | 027 | S | 0.25 | 0.25 | NA | Unknown | FALSE | FALSE | none | 2 | FST, GWAS | This study |
| JH237 | 25590 | 027 | HMR | 1 | 8 | NA | 10/8/13 | TRUE | TRUE | gyrA and nimB | 2 | FST, GWAS | This study |
| JH238 | 23025 | 027 | HMR | 1 | 8 | NA | 4/11/12 | TRUE | TRUE | gyrA and nimB | 2 | FST, GWAS | This study |
| JH239 | 23206 | 027 | HMR | 1 | 8 | NA | 5/21/12 | TRUE | TRUE | gyrA and nimB | 2 | FST, GWAS | This study |
| JH240 | 23215 | 027 | HMR | 1 | 16 | NA | 4/29/12 | TRUE | TRUE | gyrA and nimB | 2 | FST, GWAS | This study |
| JH241 | 23458 | 027 | HMR | 1 | 16 | NA | 7/2/12 | TRUE | TRUE | gyrA and nimB | 2 | FST, GWAS | This study |
| JH242 | 24568 | 027 | HMR | 1 | 16 | NA | 3/27/13 | TRUE | TRUE | gyrA and nimB | 2 | FST, GWAS | This study |
| JH243 | 24579 | 027 | HMR | 1 | 16 | NA | 4/18/13 | TRUE | TRUE | gyrA and nimB | 2 | FST, GWAS | This study |

| | | | | | | | | | | | | | |
|---|---|---|---|---|---|---|---|---|---|---|---|---|---|
| JH244 | 25605 | 027 | HMR | 1 | 8 | NA | 10/14/13 | TRUE | TRUE | gyrA and nimB | 2 | FST, GWAS | This study |
| JH245 | 26675 | 027 | HMR | 1 | 16 | NA | 4/29/14 | TRUE | TRUE | gyrA and nimB | 2 | FST, GWAS | This study |
| JH246 | 26926 | 027 | HMR | 1 | 16 | NA | 8/14/14 | TRUE | TRUE | gyrA and nimB | 2 | FST, GWAS | This study |
| JH247 | 27162 | 027 | HMR | 1 | 16 | NA | 10/3/14 | TRUE | TRUE | gyrA and nimB | 2 | FST, GWAS | This study |
| JH248 | 24391 | 027 | S | 0.25 | 0.25 | NA | 3/14/13 | TRUE | TRUE | gyrA and nimB | 2 | FST, GWAS | This study |
| JH250 | 27658 | 027 | HMR | 1 | 8 | NA | 1/7/14 | TRUE | TRUE | gyrA and nimB | 2 | FST, GWAS | This study |
| JH263 | MT1470 | 027 | HMR | 0.5 | 2 | USA | 12/22/12 | TRUE | TRUE | gyrA and nimB | 2 | FST, GWAS | This study |
| JH283 | 24921 | 027 | HMR | 0.5 | 2 | NA | NA | TRUE | TRUE | gyrA and nimB | 2 | FST, GWAS | This study |
| JH284 | 24611 | 027 | HMR | 0.25 | 2 | NA | NA | TRUE | TRUE | gyrA and nimB | 2 | FST, GWAS | This study |
| JH289 | 23038 | 027 | HMR | 0.25 | 8 | NA | 12/17/12 | TRUE | TRUE | gyrA and nimB | 2 | FST, GWAS | This study |
| JH294 | 24010 | 027 | HMR | 0.5 | 8 | NA | 1/21/13 | TRUE | TRUE | gyrA and nimB | 2 | FST, GWAS | This study |
| JH295 | 24021 | 027 | HMR | 0.25 | 2 | NA | 3/28/13 | TRUE | TRUE | gyrA and nimB | 2 | FST, GWAS | This study |
| JH298 | 24623 | 027 | HMR | 0.5 | 16 | NA | 6/8/13 | TRUE | TRUE | gyrA and nimB | 2 | FST, GWAS | This study |
| JH315 | NR-49278 | 027 | HMR | 0.25 | 2 | USA | 2011 | TRUE | TRUE | gyrA and nimB | 2 | FST, GWAS | This study |
| JH316 | NR-49279 | 027 | HMR | 0.5 | 8 | USA | 2011 | TRUE | TRUE | gyrA and nimB | 2 | FST, GWAS | This study |
| JH317 | NR-49284 | 027 | HMR | 0.5 | 8 | USA | 2011 | TRUE | TRUE | gyrA and nimB | 2 | FST, GWAS | This study |
| JH318 | NR-49286 | 027 | HMR | 0.5 | 2 | USA | 2011 | TRUE | TRUE | gyrA and nimB | 2 | FST, GWAS | This study |
| JH319 | NR-49287 | 027 | HMR | 0.5 | 2 | USA | 2010 | TRUE | TRUE | gyrA and nimB | 2 | FST, GWAS | This study |
| JH320 | NR-49288 | 027 | HMR | 0.5 | 2 | USA | 2011 | TRUE | TRUE | gyrA and nimB | 2 | FST, GWAS | This study |
| JH321 | NR-49289 | 027 | HMR | 0.25 | 2 | USA | 2010 | TRUE | TRUE | gyrA and nimB | 2 | FST, GWAS | This study |
| JH439 | L,13.7933166 | 027 | HMR | 0.25 | 2 | Germany | 2012 | TRUE | TRUE | gyrA and nimB | 2 | FST, GWAS | This study |
| JH446 | L,13.7933339 | 027 | HMR | 0.25 | 2 | Cyprus | 2013-2014 | TRUE | TRUE | gyrA and nimB | 2 | FST, GWAS | This study |
| JH476 | L,13.7933400 | 027 | HMR | 0.25 | 1 | Italy | 2012 | TRUE | TRUE | gyrA and nimB | 2 | FST, GWAS | This study |
| JH477 | L,13.7933419 | 027 | HMR | 0.25 | 4 | Italy | 2012 | TRUE | TRUE | gyrA and nimB | 2 | FST, GWAS | This study |
| JH488 | L,13.7933805 | 027 | HMR | 0.5 | 4 | UK | 2012 | TRUE | TRUE | gyrA and nimB | 2 | FST, GWAS | This study |
| JH494 | L,14.7142604 | 027 | HMR | 0.25 | 2 | Germany | 2012-2013 | TRUE | TRUE | gyrA and nimB | 2 | FST, GWAS | This study |
| JH498 | L,14.7142832 | 027 | S | 0.25 | 0.25 | Belgium | 2011-12 | FALSE | FALSE | none | 2 | FST, GWAS | This study |
| JH501 | L,14.7143010 | 027 | HMR | 0.25 | 2 | Portugal | 2013 | TRUE | TRUE | gyrA and nimB | 2 | FST, GWAS | This study |
| JH504 | L,14.7143039 | 027 | HMR | 0.5 | 4 | Poland | 2014 | TRUE | TRUE | gyrA and nimB | 2 | FST, GWAS | This study |
| JH508 | L,16.7537629 | 027 | HMR | 0.5 | 2 | Spain | 2014 | TRUE | TRUE | gyrA and nimB | 2 | FST, GWAS | This study |
| JH516 | L,15.7103779 | 027 | HMR | 0.25 | 4 | Hungary | 2014 | TRUE | TRUE | gyrA and nimB | 2 | FST, GWAS | This study |
| JH517 | L,15.7103789 | 027 | HMR | 0.25 | 4 | Hungary | ata not available 13-1 | TRUE | TRUE | gyrA and nimB | 2 | FST, GWAS | This study |
| JH539 | L,15.7104516 | 027 | HMR | 0.25 | 16 | Poland | 12/1/12 | TRUE | TRUE | gyrA and nimB | 2 | FST, GWAS | This study |
| JH540 | L,15.7104530 | 027 | HMR | 0.25 | 8 | Poland | 9/27/11 | TRUE | TRUE | gyrA and nimB | 2 | FST, GWAS | This study |
| JH541 | L,15.7104598 | 027 | HMR | 0.25 | 2 | Slovenia | 10/13/12 | TRUE | TRUE | gyrA and nimB | 2 | FST, GWAS | This study |
| JH557 | MT443 | 027 | HMR | 0.25 | 2 | USA | NA | TRUE | TRUE | gyrA and nimB | 2 | FST, GWAS | This study |
| JH558 | MT204 | 027 | HMR | 0.25 | 2 | USA | NA | TRUE | TRUE | gyrA and nimB | 2 | FST, GWAS | This study |
| JH559 | MT1300 | 027 | HMR | 0.25 | 2 | USA | NA | TRUE | TRUE | gyrA and nimB | 2 | FST, GWAS | This study |
| JH7 | 491858 | 027 | HMR | 0.5 | 8 | Israel | NA | TRUE | TRUE | gyrA and nimB | NA | Genome analysis | This study |
| JH507 | L,16.7537603 | 023 | HMR | 0.25 | 1 | Spain | 2015 | NA | NA | NA | NA | NA | NA |
| JH375 | L,13.7933429 | 023 | S | 0.25 | 0.25 | Switzerland | 2011 | FALSE | FALSE | none | 3 | FST, GWAS | This study |
| JH388 | L,14.7142515 | 023 | S | 0.25 | 0.25 | Spain | 2012-2013 | FALSE | FALSE | none | 3 | FST, GWAS | This study |
| JH409 | L,14.7142934 | 023 | S | 0.25 | 0.25 | Netherlands | 2012 | FALSE | FALSE | none | 3 | FST, GWAS | This study |
| JH412 | L,14.7143115 | 023 | S | 0.25 | 0.25 | France | 2013 | FALSE | FALSE | none | 3 | FST, GWAS | This study |
| JH425 | L,14.7143310 | 023 | S | 0.25 | 0.25 | UK | 2013 | FALSE | FALSE | none | 3 | FST, GWAS | This study |
| JH464 | L,16.7570125 | 023 | S | 0.25 | 0.25 | France | 2012 | FALSE | FALSE | none | 3 | FST, GWAS | This study |
| JH468 | L,16.7570269 | 023 | S | 0.25 | 0.25 | Ireland | 2011 | FALSE | FALSE | none | 3 | FST, GWAS | This study |
| JH532 | L,15.7104318 | 023 | S | 0.25 | 0.25 | Germany | 2014-2015 | FALSE | FALSE | none | 3 | FST, GWAS | This study |

| | | | | | | | | | | | | | |
|---|---|---|---|---|---|---|---|---|---|---|---|---|---|
| AR-1073 | AR-1073 | 020 | S | 0.25 | 0.25 | USA | 2016 | FALSE | FALSE | none | NA | Published genome | PRJNA577141 |
| AR-1096 | AR-1096 | 020 | S | 0.25 | 0.25 | USA | 2016 | FALSE | FALSE | none | NA | Published genome | PRJNA577141 |
| JH299 | 24752 | 020 | S | 0.5 | 0.25 | NA | 5/28/13 | FALSE | FALSE | none | 1 | FST, GWAS | This study |
| JH324 | NR-49300 | 020 | S | 0.5 | 0.5 | USA | 2011 | FALSE | FALSE | none | 1 | FST, GWAS | This study |
| JH325 | NR-49302 | 020 | S | 0.5 | 0.5 | USA | 2010 | FALSE | FALSE | none | 1 | FST, GWAS | This study |
| JH326 | NR-49304 | 020 | S | 1 | 0.5 | USA | 2011 | FALSE | FALSE | none | 1 | FST, GWAS | This study |
| JH360 | L,12.7903823 | 020 | S | 0.25 | 0.5 | Ireland | 2012 | FALSE | FALSE | none | 1 | FST, GWAS | This study |
| JH371 | L,13.7933304 | 020 | S | 0.25 | 0.25 | Netherlands | 2011 | FALSE | FALSE | none | 1 | FST, GWAS | This study |
| JH372 | L,13.7933283 | 020 | S | 0.25 | 0.25 | Germany | 2011 | FALSE | FALSE | none | 1 | FST, GWAS | This study |
| JH377 | L,13.7933468 | 020 | S | 0.25 | 0.25 | UK | 2012 | FALSE | FALSE | none | 1 | FST, GWAS | This study |
| JH418 | L,14.7143255 | 020 | S | 0.25 | 0.25 | Czech Rep | 2013 | FALSE | FALSE | none | 1 | FST, GWAS | This study |
| JH419 | L,14.7143236 | 020 | S | 0.25 | 0.25 | Sweden | 2014 | FALSE | FALSE | none | 1 | FST, GWAS | This study |
| JH424 | L,14.7143316 | 020 | S | 0.25 | 0.25 | Greece | 2013-2014 | FALSE | FALSE | none | 1 | FST, GWAS | This study |
| JH457 | L,16.7569720 | 020 | HMR | 0.25 | 2 | Germany | 2015-2016 | FALSE | TRUE | nimB only | 1 | FST, GWAS | This study |
| JH458 | L,16.7569753 | 020 | S | 0.25 | 0.25 | portugal | 2015 | FALSE | FALSE | none | 1 | FST, GWAS | This study |
| JH483 | L,13.7933585 | 020 | S | 0.25 | 0.25 | Poland | 2012-2013 | FALSE | FALSE | none | 1 | FST, GWAS | This study |
| JH534 | L,15.7104378 | 020 | S | 0.25 | 0.25 | France | 2015 | FALSE | FALSE | none | 1 | FST, GWAS | This study |
| AR-1075 | AR-1075 | 019 | S | 0.25 | 0.25 | USA | 2016 | FALSE | FALSE | none | NA | Published genome | PRJNA577141 |
| AR-1094 | AR-1094 | 019 | S | 0.25 | 0.25 | USA | 2016 | FALSE | FALSE | none | NA | Published genome | PRJNA577141 |
| JH322 | NR-49277 | 019 | S | 0.25 | 0.25 | USA | 2010 | FALSE | FALSE | none | 1 | FST, GWAS | This study |
| JH323 | NR-49290 | 019 | S | 1 | 1 | USA | 2011 | FALSE | FALSE | none | 2 | FST, GWAS | This study |
| JH374 | L,13.7933377 | 018 | HMR | 0.25 | 1 | Italy | 2011-2012 | TRUE | TRUE | gyrA and nimB | 1 | FST, GWAS | This study |
| JH394 | L,14.7142537 | 018 | S | 0.25 | 0.25 | Hungary | 2013 | FALSE | FALSE | none | 1 | FST, GWAS | This study |
| JH398 | L,16.7570069 | 018 | S | 0.25 | 0.25 | Ireland | 2012-2013 | FALSE | FALSE | none | 1 | FST, GWAS | This study |
| JH404 | L,14.7142825 | 018 | HMR | 0.25 | 2 | Italy | 2013 | TRUE | TRUE | gyrA and nimB | 1 | FST, GWAS | This study |
| JH405 | L,14.7142826 | 018 | HMR | 0.25 | 2 | Italy | 2013 | TRUE | TRUE | gyrA and nimB | 1 | FST, GWAS | This study |
| JH406 | L,14.7142808 | 018 | S | 0.5 | 0.25 | Italy | 2013 | TRUE | FALSE | gyrA only | 1 | FST, GWAS | This study |
| JH423 | L,14.7143303 | 018 | S | 0.25 | 0.25 | UK | 2014 | FALSE | FALSE | none | 1 | FST, GWAS | This study |
| JH431 | L,15.7104082 | 018 | S | 0.25 | 0.25 | Italy | 2012 | TRUE | FALSE | gyrA only | 1 | FST, GWAS | This study |
| JH455 | L,16.7569627 | 018 | S | 0.25 | 0.25 | Austria | 2016 | FALSE | FALSE | none | 1 | FST, GWAS | This study |
| JH461 | L,14.7142688 | 018 | S | 0.25 | 0.25 | Cyprus | 2014 | FALSE | FALSE | none | 2 | FST, GWAS | This study |
| JH485 | L,13.7933637 | 018 | S | 0.25 | 0.25 | Switzerland | 2013 | FALSE | FALSE | none | 1 | FST, GWAS | This study |
| JH213 | 24792 | 017 | HMR | 1 | 8 | NA | 6/6/13 | NA | NA | NA | NA | NA | NA |
| JH226 | 26104 | 017 | HMR | 1 | 8 | NA | 12/12/13 | NA | NA | NA | NA | NA | NA |
| JH362 | L,13.7933106 | 017 | S | 0.25 | 0.25 | Spain | 2011 | NA | NA | NA | NA | NA | NA |
| JH387 | L,14.7142505 | 017 | HMR | 0.25 | 2 | Spain | 2013-2014 | NA | NA | NA | NA | NA | NA |
| JH287 | 27118 | 017 | S | 0.25 | 0.25 | NA | 5/3/12 | TRUE | FALSE | gyrA only | 4 | FST, GWAS | This study |
| JH300 | 24778 | 017 | HMR | 0.5 | 8 | NA | 5/28/13 | TRUE | TRUE | gyrA and nimB | 4 | FST, GWAS | This study |
| JH314 | NR-49312 | 017 | S | 0.25 | 0.25 | USA | 2011 | FALSE | FALSE | none | 4 | FST, GWAS | This study |
| JH366 | L,13.7933157 | 017 | HMR | 0.25 | 2 | Germany | 2012-2013 | TRUE | TRUE | gyrA and nimB | 4 | FST, GWAS | This study |
| JH393 | L,14.7142591 | 017 | S | 0.25 | 0.25 | Germany | 2013 | TRUE | FALSE | gyrA only | 4 | FST, GWAS | This study |
| JH397 | L,14.7142654 | 017 | S | 0.25 | 0.25 | Spain | 2012-2013 | TRUE | FALSE | gyrA only | 4 | FST, GWAS | This study |
| JH417 | L,14.7143256 | 017 | S | 0.25 | 0.25 | Czech Rep | 2012-2013 | FALSE | FALSE | none | 4 | FST, GWAS | This study |
| JH427 | L,15.7103771 | 017 | S | 0.25 | 0.25 | France | 2012 | FALSE | FALSE | none | 4 | FST, GWAS | This study |
| JH429 | L,15.7103943 | 017 | S | 0.25 | 0.25 | Czech Rep | 2011 | TRUE | FALSE | gyrA only | 4 | FST, GWAS | This study |
| JH450 | L,15.7104214 | 017 | S | 0.25 | 0.25 | Italy | 2014-2015 | TRUE | FALSE | gyrA only | 4 | FST, GWAS | This study |
| JH467 | L,16.7570261 | 017 | HMR | 0.25 | 1 | Ireland | 2012 | TRUE | TRUE | gyrA and nimB | 4 | FST, GWAS | This study |
| JH482 | L,13.7933587 | 017 | S | 0.25 | 0.25 | Poland | 2013 | TRUE | FALSE | gyrA only | 4 | FST, GWAS | This study |

| | | | | | | | | | | | | | |
|---|---|---|---|---|---|---|---|---|---|---|---|---|---|
| AR-1069 | AR-1069 | 015 | S | 0.25 | 0.25 | USA | 2016 | FALSE | FALSE | none | NA | Published genome | PRJNA577141 |
| AR-1086 | AR-1086 | 015 | S | 0.25 | 0.25 | USA | 2016 | FALSE | FALSE | none | NA | Published genome | PRJNA577141 |
| JH361 | L,13.7933063 | 015 | S | 0.25 | 0.25 | Austria | 2011 | FALSE | FALSE | none | 1 | FST, GWAS | This study |
| JH384 | L,13.7933841 | 015 | S | 0.25 | 0.25 | UK | 2012-2013 | FALSE | FALSE | none | 1 | FST, GWAS | This study |
| JH385 | L,13.7933854 | 015 | S | 0.5 | 0.25 | UK | 2013 | FALSE | FALSE | none | 1 | FST, GWAS | This study |
| JH386 | L,13.7933851 | 015 | S | 0.5 | 0.25 | UK | 2013 | FALSE | FALSE | none | 1 | FST, GWAS | This study |
| JH401 | L,14.7142754 | 015 | S | 0.25 | 0.25 | Italy | 2012-2013 | FALSE | FALSE | none | 1 | FST, GWAS | This study |
| JH403 | L,14.7142815 | 015 | S | 0.25 | 0.25 | Italy | 2013-2014 | FALSE | FALSE | none | 1 | FST, GWAS | This study |
| JH408 | L,14.7142885 | 015 | S | 0.25 | 0.25 | Finland | 2013 | FALSE | FALSE | none | 1 | FST, GWAS | This study |
| JH411 | L,14.7142961 | 015 | S | 0.25 | 0.25 | Switzerland | 2013 | FALSE | FALSE | none | 1 | FST, GWAS | This study |
| JH416 | L,14.7143250 | 015 | S | 0.25 | 0.25 | Czech Rep | 2012 | FALSE | FALSE | none | 1 | FST, GWAS | This study |
| JH466 | L,16.7570203 | 015 | S | 0.25 | 0.25 | Germany | 2011 | FALSE | FALSE | none | 1 | FST, GWAS | This study |
| JH168 | MT4936 | 014-20 | HMR | 0.5 | 8 | USA | 5/3/17 | FALSE | TRUE | nimB only | 1 | FST, GWAS | This study |
| JH169 | MT4804 | 014-20 | HMR | 0.25 | 2 | USA | 12/16/16 | TRUE | TRUE | gyrA and nimB | 2 | FST, GWAS | This study |
| JH170 | SH544 | 014-20 | S | 0.5 | 0.5 | USA | 12/26/16 | FALSE | FALSE | none | 1 | FST, GWAS | This study |
| JH171 | SH553 | 014-20 | S | 0.5 | 0.25 | USA | 12/31/16 | FALSE | FALSE | none | 1 | FST, GWAS | This study |
| JH172 | SH709 | 014-20 | S | 0.25 | 0.25 | USA | 3/23/17 | FALSE | FALSE | none | 1 | FST, GWAS | This study |
| JH201 | MT1697 | 014-20 | HMR | 0.25 | 2 | USA | 3/23/13 | TRUE | TRUE | gyrA and nimB | 2 | FST, GWAS | This study |
| JH202 | MT3914 | 014-20 | HMR | 0.25 | 2 | USA | 10/11/15 | TRUE | TRUE | gyrA and nimB | 2 | FST, GWAS | This study |
| JH203 | MT3924 | 014-20 | HMR | 0.25 | 2 | USA | 10/2/15 | TRUE | TRUE | gyrA and nimB | 2 | FST, GWAS | This study |
| JH269 | SH519 | 014-20 | S | 0.5 | 0.5 | USA | 12/9/16 | TRUE | FALSE | gyrA only | 1 | FST, GWAS | This study |
| JH270 | SH512 | 014-20 | S | 0.5 | 0.25 | USA | 12/7/16 | FALSE | FALSE | none | 1 | FST, GWAS | This study |
| JH271 | SH657 | 014-20 | S | 0.5 | 0.25 | USA | 3/1/17 | FALSE | FALSE | none | 1 | FST, GWAS | This study |
| JH272 | SH660 | 014-20 | S | 0.5 | 0.5 | USA | 3/2/17 | FALSE | FALSE | none | 1 | FST, GWAS | This study |
| JH279 | MT1956 | 014-026 | S | 0.25 | 0.25 | USA | 6/7/13 | FALSE | FALSE | none | 1 | FST, GWAS | This study |
| JH391 | L,14.7142593 | 014-026 | S | 0.25 | 0.25 | Germany | 2013 | FALSE | FALSE | none | 1 | FST, GWAS | This study |
| JH192 | SL118 | 014-020 | S | 0.5 | 0.5 | NA | 7/1/11 | FALSE | FALSE | none | 1 | FST, GWAS | This study |
| JH193 | SL99 | 014-020 | S | 0.25 | 0.5 | NA | 7/1/11 | FALSE | FALSE | none | 1 | FST, GWAS | This study |
| JH194 | MT1245 | 014-020 | S | 0.5 | 1 | USA | 9/30/12 | FALSE | FALSE | none | 1 | FST, GWAS | This study |
| JH195 | MT3257 | 014-020 | S | 0.25 | 0.25 | USA | 8/13/11 | FALSE | FALSE | none | 1 | FST, GWAS | This study |
| JH196 | MT1676 | 014-020 | S | 0.25 | 0.25 | USA | 2/19/13 | FALSE | FALSE | none | 1 | FST, GWAS | This study |
| JH197 | MT1499 | 014-020 | S | 0.25 | 0.5 | USA | 11/9/12 | FALSE | FALSE | none | 1 | FST, GWAS | This study |
| JH198 | MT1294 | 014-020 | S | 0.5 | 0.25 | USA | 10/11/12 | FALSE | FALSE | none | 1 | FST, GWAS | This study |
| JH199 | MT465 | 014-020 | S | 0.25 | 0.25 | USA | 12/5/12 | FALSE | FALSE | none | 1 | FST, GWAS | This study |
| JH200 | MT3561 | 014-020 | S | 0.25 | 0.25 | USA | 6/2/15 | FALSE | FALSE | none | 1 | FST, GWAS | This study |
| JH310 | NR-49294 | 014 | S | 0.25 | 0.25 | USA | 2010 | NA | NA | NA | NA | NA | NA |
| JH551 | 23468 | 014 | HMR | 0.25 | 1 | NA | NA | NA | NA | NA | NA | NA | NA |
| JH554 | 23475 | 014 | S | 0.25 | 0.25 | NA | NA | NA | NA | NA | NA | NA | NA |
| AR-1081 | AR-1081 | 014 | S | 0.25 | 0.25 | USA | 2016 | FALSE | FALSE | none | NA | Published genome | PRJNA577141 |
| AR-1090 | AR-1090 | 014 | S | 0.25 | 0.25 | USA | 2016 | FALSE | FALSE | none | NA | Published genome | PRJNA577141 |
| AR-1091 | AR-1091 | 014 | S | 0.25 | 0.25 | USA | 2016 | FALSE | FALSE | none | NA | Published genome | PRJNA577141 |
| JH249 | 27081 | 014 | S | 1 | 0.5 | NA | 10/1/14 | FALSE | FALSE | none | 1 | FST, GWAS | This study |
| JH311 | NR-49295 | 014 | S | 0.25 | 0.25 | USA | 2011 | FALSE | FALSE | none | 1 | FST, GWAS | This study |
| JH312 | NR-49328 | 014 | S | 0.25 | 0.25 | USA | 2010 | FALSE | FALSE | none | 1 | FST, GWAS | This study |
| JH313 | NR-49329 | 014 | S | 0.25 | 0.25 | USA | 2010 | FALSE | FALSE | none | 1 | FST, GWAS | This study |
| JH369 | L,13.7933311 | 014 | S | 0.25 | 0.25 | Netherlands | 2011 | FALSE | FALSE | none | 1 | FST, GWAS | This study |
| JH370 | L,13.7933281 | 014 | S | 0.25 | 0.25 | Germany | 2011 | TRUE | FALSE | gyrA only | 1 | FST, GWAS | This study |
| JH373 | L,13.7933432 | 014 | S | 0.25 | 0.25 | Switzerland | 2012 | FALSE | FALSE | none | 1 | FST, GWAS | This study |

| | | | | | | | | | | | | | |
|---|---|---|---|---|---|---|---|---|---|---|---|---|---|
| JH376 | L,13.7933458 | 014 | S | 0.25 | 0.25 | UK | 2011 | FALSE | FALSE | none | 1 | FST, GWAS | This study |
| JH392 | L,14.7142559 | 014 | S | 0.25 | 0.25 | Greece | 2014-2015 | FALSE | FALSE | none | 1 | FST, GWAS | This study |
| JH400 | L,14.7142740 | 014 | S | 0.25 | 0.25 | Italy | 2012-2013 | FALSE | FALSE | none | 1 | FST, GWAS | This study |
| JH428 | L,15.7103888 | 014 | S | 0.25 | 0.25 | Sweden. | 2012 | FALSE | FALSE | none | 1 | FST, GWAS | This study |
| JH430 | L,15.7104142 | 014 | S | 0.25 | 0.25 | Ireland | 2012 | FALSE | FALSE | none | 1 | FST, GWAS | This study |
| JH432 | L,15.7104045 | 014 | S | 0.25 | 0.25 | UK | 2012 | FALSE | FALSE | none | 1 | FST, GWAS | This study |
| ATCC43596 | ATCC43596 | 012 | S | 0.25 | 0.25 | NA | NA | NA | NA | NA | NA | NA | NA |
| CD630 | CD630 | 012 | S | 0.5 | 0.5 | Switzerland | 1982 | FALSE | FALSE | none | none | Genome analysis | PRJNA78 |
| JH167 | MT5065 | 012 | S | 0.25 | 0.25 | USA | 5/2/17 | FALSE | FALSE | none | 1 | FST, GWAS | This study |
| JH365 | L,13.7933184 | 012 | HMR | 0.25 | 1 | Czech Rep | 2011 | FALSE | FALSE | none | 1 | FST, GWAS | This study |
| JH367 | L,13.7933262 | 012 | S | 0.25 | 0.25 | Italy | 2011 | FALSE | FALSE | none | 1 | FST, GWAS | This study |
| JH421 | L,14.7143267 | 012 | S | 0.25 | 0.25 | Slovenia | 2013 | FALSE | FALSE | none | 1 | FST, GWAS | This study |
| JH422 | L,14.7143317 | 012 | S | 0.25 | 0.25 | Greece | 2013 | FALSE | FALSE | none | 1 | FST, GWAS | This study |
| JH452 | L,15.7104335 | 012 | S | 0.25 | 0.25 | Germany | 2014-2015 | FALSE | FALSE | none | 1 | FST, GWAS | This study |
| JH454 | L,16.7569633 | 012 | S | 0.25 | 0.25 | Austria | 2013 | FALSE | FALSE | none | 1 | FST, GWAS | This study |
| JH456 | L,16.7569767 | 012 | S | 0.25 | 0.25 | portugal | 2015-2016 | FALSE | FALSE | none | 1 | FST, GWAS | This study |
| JH470 | L,16.7537614 | 012 | S | 0.25 | 0.5 | Spain | 2012 | FALSE | FALSE | none | 1 | FST, GWAS | This study |
| IT1001 | IT1001 | 010 | HMR | 0.5 | 8 | Italy | NA | FALSE | FALSE | none | NA | Genome analysis | This study |
| IT1002 | IT1002 | 010 | HMR | 0.5 | 8 | Italy | NA | FALSE | FALSE | none | NA | Genome analysis | This study |
| JH395 | L,14.7142616 | 010 | S | 0.25 | 0.25 | Austria | 2013 | FALSE | FALSE | none | 1 | FST, GWAS | This study |
| JH420 | L,14.7143300 | 010 | S | 0.25 | 0.25 | UK | 2014 | FALSE | FALSE | none | 1 | FST, GWAS | This study |
| JH533 | L,15.7104380 | 010 | HMR | 0.25 | 8 | France | 2014-2015 | TRUE | FALSE | gyrA only | 1 | FST, GWAS | This study |
| JH364 | L,13.7933155 | 005 | HMR | 0.25 | 2 | Germany | 2012-2013 | TRUE | TRUE | gyrA and nimB | 2 | FST, GWAS | This study |
| JH382 | L,13.7933696 | 005 | S | 0.25 | 0.25 | UK | 2013-2014 | FALSE | FALSE | none | 1 | FST, GWAS | This study |
| JH383 | L,13.7933773 | 005 | S | 0.25 | 0.25 | France | 2012 | FALSE | FALSE | none | 1 | FST, GWAS | This study |
| JH396 | L,14.7142622 | 005 | S | 0.25 | 0.25 | Austria | 2013 | FALSE | FALSE | none | 1 | FST, GWAS | This study |
| JH402 | L,14.7142854 | 005 | S | 0.25 | 0.25 | Belgium | 2013-2014 | FALSE | FALSE | none | 1 | FST, GWAS | This study |
| JH407 | L,14.7142893 | 005 | S | 0.25 | 0.25 | Finland | 2013-2014 | FALSE | FALSE | none | 1 | FST, GWAS | This study |
| JH415 | L,14.7143227 | 005 | S | 0.25 | 0.25 | Sweden | 2012-2013 | FALSE | FALSE | none | 1 | FST, GWAS | This study |
| JH426 | L,15.7103831 | 005 | S | 0.25 | 0.25 | Hungary | 2014 | FALSE | FALSE | none | 1 | FST, GWAS | This study |
| JH449 | L,15.7104188 | 005 | S | 0.25 | 0.25 | Netherlands. | 2015 | FALSE | FALSE | none | 1 | FST, GWAS | This study |
| JH453 | L,15.7104528 | 005 | S | 0.25 | 0.25 | Poland | 2014-2015 | FALSE | FALSE | none | 1 | FST, GWAS | This study |
| JH469 | L,16.7537597 | 005 | S | 0.25 | 0.25 | Spain | 2011 | FALSE | FALSE | none | 1 | FST, GWAS | This study |
| JH531 | L,15.7104232 | 005 | S | 0.25 | 0.25 | Cyprus | 2014-2015 | FALSE | FALSE | none | 1 | FST, GWAS | This study |
| JH381 | L,13.7933706 | 002 | S | 0.25 | 0.25 | France | 2012 | NA | NA | NA | NA | NA | NA |
| JH410 | L,14.7142968 | 002 | S | 0.25 | 0.25 | Switzerland | 2013 | NA | NA | NA | NA | NA | NA |
| JH462 | L,16.7570111 | 002 | S | 0.25 | 0.25 | France | 2015 | NA | NA | NA | NA | NA | NA |
| AR-1070 | AR-1070 | 002 | S | 0.25 | 0.25 | USA | 2016 | FALSE | FALSE | none | NA | Published genome | PRJNA577141 |
| AR-1074 | AR-1074 | 002 | S | 0.25 | 0.25 | USA | 2016 | TRUE | FALSE | gyrA only | NA | Published genome | PRJNA577141 |
| AR-1084 | AR-1084 | 002 | S | 0.25 | 0.25 | USA | 2016 | FALSE | FALSE | none | NA | Published genome | PRJNA577141 |
| JH163 | MT4941 | 002 | S | 0.25 | 0.25 | USA | 2/23/17 | FALSE | FALSE | none | 2 | FST, GWAS | This study |
| JH164 | MT5006 | 002 | S | 0.25 | 0.25 | USA | 3/31/17 | FALSE | FALSE | none | 1 | FST, GWAS | This study |
| JH165 | MT4961 | 002 | S | 0.25 | 0.25 | USA | 2/28/17 | FALSE | FALSE | none | 1 | FST, GWAS | This study |
| JH166 | SH540 | 002 | S | 0.25 | 0.25 | USA | 12/25/16 | FALSE | FALSE | none | 1 | FST, GWAS | This study |
| JH273 | SH650 | 002 | S | 0.5 | 0.5 | USA | 2/26/17 | FALSE | FALSE | none | 1 | FST, GWAS | This study |
| JH306 | NR-49305 | 002 | S | 0.5 | 0.5 | USA | 2011 | FALSE | FALSE | none | 1 | FST, GWAS | This study |
| JH307 | NR-49306 | 002 | S | 0.5 | 0.5 | USA | 2011 | FALSE | FALSE | none | 1 | FST, GWAS | This study |
| JH308 | NR-49307 | 002 | S | 0.5 | 0.5 | USA | 2011 | FALSE | FALSE | none | 1 | FST, GWAS | This study |

| | | | | | | | | | | | | | |
|---|---|---|---|---|---|---|---|---|---|---|---|---|---|
| JH309 | NR-49308 | 002 | S | 0.5 | 0.5 | USA | 2011 | FALSE | FALSE | none | 1 | FST, GWAS | This study |
| JH359 | L,12.7903758 | 002 | S | 0.25 | 0.25 | France | 2012 | FALSE | FALSE | none | 1 | FST, GWAS | This study |
| JH368 | L,13.7933282 | 002 | S | 0.25 | 0.25 | Germany | 2012-2013 | FALSE | FALSE | none | 1 | FST, GWAS | This study |
| JH378 | L,13.7933590 | 002 | S | 0.25 | 0.25 | Poland | 2012 | FALSE | FALSE | none | 1 | FST, GWAS | This study |
| JH390 | L,14.7142552 | 002 | S | 0.25 | 0.25 | Hungary | 2013 | FALSE | FALSE | none | 1 | FST, GWAS | This study |
| JH399 | L,14.7142738 | 002 | S | 0.25 | 0.25 | Italy | 2012-2013 | FALSE | FALSE | none | 1 | FST, GWAS | This study |
| JH414 | L,14.7143199 | 002 | S | 0.25 | 0.25 | Denmark | 2012-2013 | FALSE | FALSE | none | 1 | FST, GWAS | This study |
| JH459 | L,16.7570007 | 002 | S | 0.25 | 0.25 | SloveniaÂ· | 2015-2016 | FALSE | FALSE | none | 1 | FST, GWAS | This study |
| JH460 | L,16.7570014 | 002 | S | 0.25 | 0.25 | Slovenia | 2015-2016 | FALSE | FALSE | none | 1 | FST, GWAS | This study |
| JH304 | NR-49292 | 001-072 | S | 0.5 | 0.5 | USA | 2011 | NA | NA | NA | NA | NA | NA |
| JH305 | NR-49293 | 001-072 | S | 0.5 | 0.5 | USA | 2011 | FALSE | FALSE | none | 1 | FST, GWAS | This study |
| 17/27 | 17/27 | 001 | HMR | 0.5 | 16 | NA | NA | TRUE | FALSE | gyrA only | NA | Genome analysis | This study |
| 7/34 | 7/34 | 001 | HMR | 0.5 | 16 | NA | NA | TRUE | FALSE | gyrA only | NA | Genome analysis | This study |
| JH161 | MT5000 | 001 | S | 0.25 | 0.25 | USA | 3/30/17 | FALSE | FALSE | none | 1 | FST, GWAS | This study |
| JH162 | SH564 | 001 | S | 0.25 | 0.25 | USA | 1/12/17 | TRUE | FALSE | gyrA only | 1 | FST, GWAS | This study |
| JH214 | 25035 | 001 | HMR | 1 | 8 | NA | 7/10/13 | TRUE | TRUE | gyrA and nimB | 1 | FST, GWAS | This study |
| JH215 | 25036 | 001 | HMR | 1 | 8 | NA | 7/25/13 | TRUE | TRUE | gyrA and nimB | 1 | FST, GWAS | This study |
| JH217 | 25353 | 001 | HMR | 1 | 8 | NA | 7/10/13 | TRUE | TRUE | gyrA and nimB | 1 | FST, GWAS | This study |
| JH218 | 25365 | 001 | HMR | 1 | 8 | NA | 8/8/13 | TRUE | TRUE | gyrA and nimB | 1 | FST, GWAS | This study |
| JH219 | 25371 | 001 | HMR | 1 | 8 | NA | 6/4/13 | TRUE | TRUE | gyrA and nimB | 1 | FST, GWAS | This study |
| JH220 | 25578 | 001 | HMR | 1 | 8 | NA | 9/2/13 | TRUE | TRUE | gyrA and nimB | 1 | FST, GWAS | This study |
| JH224 | 25641 | 001 | HMR | 1 | 8 | NA | 10/14/13 | TRUE | TRUE | gyrA and nimB | 1 | FST, GWAS | This study |
| JH227 | 26118 | 001 | HMR | 1 | 8 | NA | 2/5/14 | TRUE | TRUE | gyrA and nimB | 1 | FST, GWAS | This study |
| JH228 | 26128 | 001 | HMR | 1 | 8 | NA | 1/31/14 | TRUE | TRUE | gyrA and nimB | 1 | FST, GWAS | This study |
| JH274 | MT4802 | 001 | S | 0.5 | 0.25 | USA | 12/14/16 | TRUE | FALSE | gyrA only | 1 | FST, GWAS | This study |
| JH363 | L,13.7933145 | 001 | S | 0.25 | 0.5 | Spain | 2011-2012 | FALSE | FALSE | none | 1 | FST, GWAS | This study |
| JH379 | L,13.7933623 | 001 | HMR | 0.25 | 2 | Switzerland | 2013 | TRUE | TRUE | gyrA and nimB | 1 | FST, GWAS | This study |
| JH380 | L,13.7933657 | 001 | HMR | 0.25 | 2 | Latvia | 2012 | TRUE | TRUE | gyrA and nimB | 1 | FST, GWAS | This study |
| JH389 | L,14.7142611 | 001 | HMR | 0.25 | 2 | Austria | 2012 | TRUE | TRUE | gyrA and nimB | 1 | FST, GWAS | This study |
| JH413 | L,14.7143170 | 001 | S | 0.25 | 0.25 | Ireland | 2012 | FALSE | FALSE | none | 1 | FST, GWAS | This study |
| JH433 | L,15.7104181 | 001 | S | 0.25 | 0.25 | Netherlands | 2012 | FALSE | FALSE | none | 1 | FST, GWAS | This study |
| JH443 | L,13.7933151 | 001 | HMR | 0.25 | 2 | Germany | 2014 | TRUE | TRUE | gyrA and nimB | 1 | FST, GWAS | This study |
| JH448 | L,13.7933296 | 001 | HMR | 0.25 | 8 | Germany | 2014-2015 | TRUE | TRUE | gyrA and nimB | 1 | FST, GWAS | This study |
| JH451 | L,15.7104308 | 001 | HMR | 0.25 | 1 | Bulgaria | 2015 | TRUE | FALSE | gyrA only | 1 | FST, GWAS | This study |
| JH463 | L,16.7570167 | 001 | HMR | 0.25 | 2 | Czech Rep | 2015 | TRUE | TRUE | gyrA and nimB | 1 | FST, GWAS | This study |
| JH465 | L,16.7570255 | 001 | S | 0.5 | 0.25 | Ireland | 2011 | FALSE | FALSE | none | 1 | FST, GWAS | This study |
| JH301 | 26168 | 404 | S | 0.25 | 0.25 | NA | 2011 | FALSE | FALSE | none | 1 | FST, GWAS | This study |
| JH475 | L,13.7933408 | 356 | S | 0.25 | 0.25 | Italy | 2011 | NA | NA | NA | NA | NA | NA |
| JH496 | L,14.7142751 | 356 | HMR | 0.25 | 1 | Italy | 2013 | NA | NA | NA | NA | NA | NA |
| JH444 | L,13.7933258 | 356 | HMR | 0.25 | 2 | Italy | 2014 | TRUE | TRUE | gyrA and nimB | 1 | FST, GWAS | This study |
| JH445 | L,13.7933268 | 356 | S | 0.25 | 0.25 | Italy | 2014 | TRUE | FALSE | gyrA only | 1 | FST, GWAS | This study |
| JH474 | L,13.7933406 | 356 | S | 0.25 | 0.25 | Italy | 2011-2012 | TRUE | FALSE | gyrA only | 1 | FST, GWAS | This study |
| JH484 | L,13.7933605 | 356 | S | 0.25 | 0.25 | Italy | 2012-2013 | TRUE | FALSE | gyrA only | 1 | FST, GWAS | This study |
| JH497 | L,14.7142749 | 356 | HMR | 0.25 | 2 | Italy | 2015 | TRUE | TRUE | gyrA and nimB | 1 | FST, GWAS | This study |
| JH499 | L,14.7142776 | 356 | S | 0.25 | 0.25 | Italy | 2011 | TRUE | FALSE | gyrA only | 1 | FST, GWAS | This study |
| JH500 | L,14.7142812 | 356 | S | 0.25 | 0.25 | Italy | 2013 | TRUE | FALSE | gyrA only | 1 | FST, GWAS | This study |
| JH528 | L,15.7104072 | 356 | S | 0.25 | 0.25 | Italy | 2013-2014 | TRUE | FALSE | gyrA only | 1 | FST, GWAS | This study |
| JH293 | 23828 | 255 | S | 0.5 | 0.5 | NA | 11/30/12 | FALSE | FALSE | none | 1 | FST, GWAS | This study |

| | | | | | | | | | | | | | |
|---|---|---|---|---|---|---|---|---|---|---|---|---|---|
| JH223 | 25640 | 231 | HMR | 0.25 | 1 | NA | 10/9/13 | TRUE | FALSE | gyrA only | 1 | FST, GWAS | This study |
| JH216 | 25069 | 198 | HMR | 1 | 16 | NA | 8/16/13 | TRUE | TRUE | gyrA and nimB | 2 | FST, GWAS | This study |
| JH225 | 25913 | 198 | HMR | 1 | 8 | NA | 11/12/13 | TRUE | TRUE | gyrA and nimB | 2 | FST, GWAS | This study |
| JH231 | 26266 | 198 | HMR | 1 | 8 | NA | 2/11/14 | TRUE | TRUE | gyrA and nimB | 2 | FST, GWAS | This study |
| JH232 | 26403 | 198 | HMR | 1 | 8 | NA | 3/19/14 | TRUE | TRUE | gyrA and nimB | 2 | FST, GWAS | This study |
| JH234 | 26741 | 198 | HMR | 1 | 8 | NA | 6/2/14 | TRUE | TRUE | gyrA and nimB | 2 | FST, GWAS | This study |
| JH236 | 27547 | 198 | HMR | 1 | 8 | NA | 9/16/14 | TRUE | TRUE | gyrA and nimB | 2 | FST, GWAS | This study |
| JH435 | L,12.7903856 | 198 | HMR | 0.25 | 2 | Hungary | 2011 | TRUE | TRUE | gyrA and nimB | 2 | FST, GWAS | This study |
| JH436 | L,12.7903874 | 198 | HMR | 0.25 | 2 | Hungary | 2012 | TRUE | TRUE | gyrA and nimB | 2 | FST, GWAS | This study |
| JH437 | L,12.7903855 | 198 | HMR | 0.25 | 2 | Hungary | 2011-2012 | TRUE | TRUE | gyrA and nimB | 2 | FST, GWAS | This study |
| JH492 | L,14.7142538 | 198 | HMR | 0.25 | 2 | Hungary | 2012-2013 | TRUE | TRUE | gyrA and nimB | 2 | FST, GWAS | This study |
| JH493 | L,14.7142532 | 198 | HMR | 0.25 | 2 | Hungary | 2012 | TRUE | TRUE | gyrA and nimB | 2 | FST, GWAS | This study |
| JH513 | L,15.7103788 | 198 | HMR | 0.25 | 2 | Hungary | 2013 | TRUE | TRUE | gyrA and nimB | 2 | FST, GWAS | This study |
| JH514 | L,15.7103776 | 198 | HMR | 0.25 | 2 | Hungary | 2014 | TRUE | TRUE | gyrA and nimB | 2 | FST, GWAS | This study |
| JH515 | L,15.7103794 | 198 | HMR | 0.25 | 2 | Hungary | 2014 | TRUE | TRUE | gyrA and nimB | 2 | FST, GWAS | This study |
| JH542 | L,16.7569662 | 198 | HMR | 0.25 | 2 | Hungary | NA | TRUE | TRUE | gyrA and nimB | 2 | FST, GWAS | This study |
| JH543 | L,16.7569653 | 198 | HMR | 0.25 | 2 | Hungary | NA | TRUE | TRUE | gyrA and nimB | 2 | FST, GWAS | This study |
| JH544 | L,16.7569675 | 198 | HMR | 0.25 | 8 | Hungary | NA | TRUE | TRUE | gyrA and nimB | 2 | FST, GWAS | This study |
| JH229 | 26137 | 176 | HMR | 1 | 8 | NA | 1/22/14 | TRUE | TRUE | gyrA and nimB | 2 | FST, GWAS | This study |
| JH230 | 26138 | 176 | HMR | 1 | 8 | NA | 2/5/14 | TRUE | TRUE | gyrA and nimB | 2 | FST, GWAS | This study |
| JH233 | 26668 | 176 | HMR | 1 | 16 | NA | 5/20/14 | TRUE | TRUE | gyrA and nimB | 2 | FST, GWAS | This study |
| JH235 | 27098 | 176 | HMR | 1 | 16 | NA | 9/2/14 | TRUE | TRUE | gyrA and nimB | 2 | FST, GWAS | This study |
| JH442 | L,13.7933192 | 176 | HMR | 0.25 | 2 | Czech Rep | 2014 | TRUE | TRUE | gyrA and nimB | 2 | FST, GWAS | This study |
| JH502 | L,14.7143044 | 176 | HMR | 0.25 | 2 | Poland | 2014 | TRUE | TRUE | gyrA and nimB | 2 | FST, GWAS | This study |
| JH503 | L,14.7143027 | 176 | HMR | 0.25 | 2 | Portugal | 2014 | TRUE | TRUE | gyrA and nimB | 2 | FST, GWAS | This study |
| JH506 | L,14.7143132 | 176 | HMR | 0.25 | 1 | Germany | 2014 | TRUE | TRUE | gyrA and nimB | 2 | FST, GWAS | This study |
| JH509 | L,13.7933591 | 176 | HMR | 0.25 | 2 | Poland | 2013 | TRUE | TRUE | gyrA and nimB | 2 | FST, GWAS | This study |
| JH511 | L,14.7143134 | 176 | HMR | 0.25 | 2 | Germanv | 2013 | TRUE | TRUE | gyrA and nimB | 2 | FST, GWAS | This study |
| JH519 | L,15.7103958 | 176 | HMR | 0.25 | 2 | Czech Rep | 2013 | TRUE | TRUE | gyrA and nimB | 2 | FST, GWAS | This study |
| JH520 | L,15.7103953 | 176 | HMR | 0.25 | 2 | Czech Rep | 2014 | TRUE | TRUE | gyrA and nimB | 2 | FST, GWAS | This study |
| JH521 | L,15.7103954 | 176 | HMR | 0.25 | 4 | Czech Rep | 2013 | TRUE | TRUE | gyrA and nimB | 2 | FST, GWAS | This study |
| JH522 | L,15.7103945 | 176 | HMR | 0.25 | 2 | Czech Rep | 2014 | TRUE | TRUE | gyrA and nimB | 2 | FST, GWAS | This study |
| JH523 | L,15.7103947 | 176 | HMR | 0.25 | 2 | Czech Rep | 2014 | TRUE | TRUE | gyrA and nimB | 2 | FST, GWAS | This study |
| JH526 | L,15.7104058 | 176 | HMR | 0.25 | 2 | UK | 2014 | TRUE | TRUE | gyrA and nimB | 2 | FST, GWAS | This study |
| JH527 | L,15.7104059 | 176 | HMR | 0.25 | 2 | UK | 2013-2014 | TRUE | TRUE | gyrA and nimB | 2 | FST, GWAS | This study |
| JH538 | L,15.7104526 | 176 | HMR | 0.25 | 2 | Poland | 2015 | TRUE | TRUE | gyrA and nimB | 2 | FST, GWAS | This study |
| JH548 | L,16.7570163 | 176 | HMR | 0.25 | 2 | Czech Rep. | NA | TRUE | TRUE | gyrA and nimB | 2 | FST, GWAS | This study |
| JH180 | SH1073 | 137 | S | 0.25 | 0.25 | USA | 12/9/17 | FALSE | FALSE | none | 1 | FST, GWAS | This study |
| JH211 | 24531 | 137 | HMR | 0.25 | 1 | NA | 3/4/13 | TRUE | FALSE | gyrA only | 1 | FST, GWAS | This study |
| JH221 | 25603 | 137 | HMR | 0.5 | 2 | NA | 10/7/13 | TRUE | FALSE | gyrA only | 1 | FST, GWAS | This study |
| JH332 | NR-49324 | 126 | S | 0.5 | 0.5 | USA | 2012 | FALSE | FALSE | none | 1 | FST, GWAS | This study |
| JH441 | L,13.7933143 | 126 | S | 0.25 | 0.25 | Spain | 2011-2012 | TRUE | FALSE | gyrA only | 5 | FST, GWAS | This study |
| JH481 | L,13.7933525 | 126 | S | 0.25 | 0.25 | Portugal | 2013-2014 | TRUE | FALSE | gyrA only | 5 | FST, GWAS | This study |
| JH491 | L,14.7142556 | 126 | S | 0.25 | 0.25 | Greece | 2012 | TRUE | FALSE | gyrA only | 5 | FST, GWAS | This study |
| JH495 | L,14.7142741 | 126 | S | 0.25 | 0.25 | Italy | 2013 | TRUE | FALSE | gyrA only | 5 | FST, GWAS | This study |
| JH505 | L,14.7143118 | 126 | S | 0.25 | 0.25 | Germany | 2014 | FALSE | FALSE | none | 5 | FST, GWAS | This study |
| JH512 | L,14.7143235 | 126 | S | 0.25 | 0.25 | Sweden | 2013 | FALSE | FALSE | none | 5 | FST, GWAS | This study |
| JH518 | L,15.7103850 | 126 | S | 0.25 | 0.25 | Greece | 2014 | FALSE | FALSE | none | 5 | FST, GWAS | This study |

| | | | | | | | | | | | | | |
|---|---|---|---|---|---|---|---|---|---|---|---|---|---|
| JH525 | L,15.7104057 | 126 | S | 0.25 | 0.25 | UK | 2014 | TRUE | FALSE | gyrA only | 5 | FST, GWAS | This study |
| JH537 | L,15.7104489 | 126 | S | 0.25 | 0.25 | Spain | 2016 | TRUE | FALSE | gyrA only | 5 | FST, GWAS | This study |
| JH549 | L,16.7570209 | 126 | S | 0.25 | 0.25 | Germany | NA | TRUE | FALSE | gyrA only | 5 | FST, GWAS | This study |
| AR-1078 | AR-1078 | 106 | S | 0.25 | 0.25 | USA | 2016 | FALSE | FALSE | none | NA | Published genome | PRJNA577141 |
| AR-1085 | AR-1085 | 106 | S | 0.25 | 0.25 | USA | 2016 | FALSE | FALSE | none | NA | Published genome | PRJNA577141 |
| AR-1087 | AR-1087 | 106 | S | 0.25 | 0.25 | USA | 2016 | FALSE | FALSE | none | NA | Published genome | PRJNA577141 |
| AR-1089 | AR-1089 | 106 | S | 0.25 | 0.25 | USA | 2016 | FALSE | FALSE | none | NA | Published genome | PRJNA577141 |
| AR-1093 | AR-1093 | 106 | S | 0.25 | 0.25 | USA | 2016 | FALSE | FALSE | none | NA | Published genome | PRJNA577141 |
| 71/01 | 71/01 | 106 | HMR | 0.5 | 8 | NA | NA | TRUE | TRUE | gyrA and nimB | NA | Genome analysis | This study |
| 72/25 | 72/25 | 106 | HMR | 0.5 | 16 | NA | NA | TRUE | TRUE | gyrA and nimB | NA | Genome analysis | This study |
| JH222 | 25626 | 106 | HMR | 1 | 8 | NA | 10/24/13 | TRUE | TRUE | gyrA and nimB | 2 | FST, GWAS | This study |
| JH281 | 24496 | 106 | S | 0.25 | 0.25 | NA | NA | FALSE | FALSE | none | 1 | FST, GWAS | This study |
| JH331 | NR-49321 | 106 | S | 0.5 | 0.5 | USA | 2012 | FALSE | FALSE | none | 1 | FST, GWAS | This study |
| JH447 | L,13.7933329 | 106 | HMR | 0.5 | 2 | Cyprus | 2014-2015 | TRUE | TRUE | gyrA and nimB | 1 | FST, GWAS | This study |
| JH471 | L,16.7537658 | 106 | S | 0.25 | 0.25 | Spain | 2011 | FALSE | FALSE | none | 1 | FST, GWAS | This study |
| JH473 | L,13.7933403 | 106 | S | 0.25 | 0.25 | Italy | 2011-2012 | FALSE | FALSE | none | 1 | FST, GWAS | This study |
| JH480 | L,13.7933455 | 106 | HMR | 0.25 | 8 | UK | 2012 | TRUE | TRUE | gyrA and nimB | 1 | FST, GWAS | This study |
| JH486 | L,13.7933691 | 106 | HMR | 0.25 | 2 | UK | 2013 | TRUE | TRUE | gyrA and nimB | 1 | FST, GWAS | This study |
| JH487 | L,13.7933707 | 106 | S | 0.25 | 0.25 | France | 2013 | FALSE | FALSE | none | 1 | FST, GWAS | This study |
| JH490 | L,14.7142513 | 106 | S | 0.25 | 0.25 | Spain | 2012-2013 | TRUE | FALSE | gyrA only | 1 | FST, GWAS | This study |
| JH524 | L,15.7104005 | 106 | S | 0.25 | 0.25 | UK | 2014 | FALSE | FALSE | none | 1 | FST, GWAS | This study |
| JH529 | L,15.7104014 | 106 | HMR | 0.25 | 8 | UK | 2012-2014 | TRUE | TRUE | gyrA and nimB | 1 | FST, GWAS | This study |
| JH530 | L,15.7104213 | 106 | S | 0.25 | 0.25 | Italy | 2013-2014 | FALSE | FALSE | none | 1 | FST, GWAS | This study |
| JH535 | L,15.7104503 | 106 | HMR | 0.25 | 2 | Spain | 2014 | TRUE | TRUE | gyrA and nimB | 1 | FST, GWAS | This study |
| JH536 | L,15.7104487 | 106 | S | 0.25 | 0.25 | Spain | 2014-2015 | TRUE | FALSE | gyrA only | 1 | FST, GWAS | This study |

**Supplementary Table 2.** Primers used in this study.

| Primers | Note | Sequence 5′ – 3′ |
|---|---|---|
| nimB_SacI_F | | taagcagagctcagaaaggatgatattatgtttaaagaaatgagattgaaaaaaagaga |
| nimB_BamHI_R | pRPF185-*nimB* | tgcttaggatccttatatacgtgaagctttaccagaaatatggtcaat |
| nimB_prom_KpnI_F2 | | taagcaggtacccacttactgttaagccacctagatac |
| nimB_BamHI_R2 | pRPF185-P*nimB* (Promoter nimB+nimB) | tgcttaggatccttatatacgtgaagctttaccagaaatatg |
| PnimOnly_NheI_1728_F2 | | taagcagctagccacttactgttaagccacct |
| PnimOnly_SacI_1728_R2 | P*nimB* ::mcherryOpt (Promoter nimB only) | tgcttagagctcaatatcatcctttctaattttgatgct |
| nimB_BamHI_613_F2 | | taagcaggatccatgtttaaagaaatgagattgaaaaaaagagaaatgacaaaagaaga |
| nimB_XhoI_613_R2 | pWL613a-NimB | tgcttactcgagttatatacgtgaagctttaccagaaatatggtcaatttctat |
| | **qPCR** | |
| mcherry_qpcr_F | | tgggatgggaagcatcatctg |
| mcherry_qpcr_R | *C. difficile* codon optimized mcherry | acttcagcatcataatgacctcca |
| rbsA_qpcr_F | | agaatttcctggtgtcagagca |
| rbsA_qpcr_R | *rbsA* | agaagttttcgaatgtgcaccag |
| vanG_qpcr_F | | gtgcgtgcaggttcatcatt |
| vanG_qpcr_R | *vanG* | actgcacaccctacctcaaa |
| CDR20291_1226_qpcr_F | | caagcaattggtggtggtgg |
| CDR20291_1226_qpcr_R | *hatT* | ggcaataccatatgttgcccc |
| CDR20291_0780_qpcr_F | | aagagatggtggagatgtagagc |
| CDR20291_0780_qpcr_R | *CDR20291_0780* | catagtcgctcctggacacc |
| CDR20291_0782_qpcr_F | | tgtggcaaattcaaatcctgttgt |
| CDR20291_0782_qpcr_R | *hsmR* | gctaaaggtacggtttcaggc |
| CDR20291_1472_qpcr_F | | ggggtgttattgcagttgacc |
| CDR20291_1472_qpcr_R | *CDR20291_1472* | gcacctccacaatcttctgc |
| CDR20291_1647_qpcr_F | | caacaagtggtcttgacccg |
| CDR20291_1647_qpcr_R | *CDR20291_1647* | ctgtcacaaagctccgttgc |
| def1_qpcr_F | | gctgcatgccaagtaggagt |
| def1_qpcr_R | *def1* | ttaccccatacttctggaaaactt |
| CDR20291_1503_qpcr_F | | cccatacgatttggagtttcagc |
| CDR20291_1503_qpcr_R | *CDR20291_1503* | acggtgcaccattcttttgc |
| CDR20291_1652_qpcr_F | | agaagaaaaagcaacatgtggagg |
| CDR20291_1652_qpcr_R | *CDR20291_1652* | catgatatttccagaccacaattca |
| ribD_qpcr_F | | acaaggaggcgttttcccat |
| ribD_qpcr_R | *ribD* | tggctctaatcatgctgaagt |
| dinR_qpcr_F | | gagctaaaatgggttctccagc |
| dinR_qpcr_R | *dinR* | aggaccctactaaaccaagagc |
| oppA_qpcr_F | | cagcaggtgagagtgaccaa |
| oppA_qpcr_R | *oppA* | tgagcatcttgattagcccaa |
| CDR20291_3199_qpcr_F | | tgtctctctgggctctctgg |
| CDR20291_3199_qpcr_R | *CDR20291_3199* | ttggcaagtgtttgggtgtg |
| CDR20291_0395_qpcr_F | | ggagggaagctgcaatgtct |
| CDR20291_0395_qpcr_R | *CDR20291_0395* | tgtgaaagttctatgcaggtga |
| CDR20291_1650_qpcr_F | | tgaaccttcaatatggttggca |

| | | |
|---|---|---|
| CDR20291_1650_qpcr_R | *CDR20291_1650* | tctcccctacttatctgtgctct |
| CDR20291_1553_qpcr_F | | agcagacccttggaatgcag |
| CDR20291_1553_qpcr_R | *CDR20291_1553* | agctgtgtccctttttgctga |
| purH_qpcr_F | | acaggagctgttttagcgtca |
| purH_qpcr_R | *purH* | agccacctggttgaactactg |
| CDR20291_1647_qpcr_F | | caacaagtggtcttgacccg |
| CDR20291_1647 _qpcr_R | *CDR20291_1647* | ctgtcacaaagctccgttgc |
| CDR20291_1649_qpcr_F | | caacaagtggtcttgacccg |
| CDR20291_1649_qpcr_R | *CDR20291_1649* | ctgtcacaaagctccgttgc |
| nrdF_qpcr_F | | tcatgtgttttggtttcagtgct |
| nrdF_qpcr_R | *nrdF* | tgggcttgaaaaagaggttgtg |
| dacF_qpcr_F | | tgcttgtgttgctatggctg |
| dacF_qpcr_R | *dacF* | agtggtttgcaactggaagtc |
| groES_qpcr_F | | accaggagcagctaaagagc |
| groES_qpcr_R | *groES* | ccttatctcccactgtcaattcca |
| CDR20291_0804_qpcr_F | | ccctgtaacaagaccagcagt |
| CDR20291_0804_qpcr_R | *CDR20291_0804* | gacatgtctgctgatggtagc |
| leuS_qpcr_F | | accacatctgctaactcgtca |
| leuS_qpcr_R | *leuS* | ttgcagcaccaccagaaaga |
| CDR20291_2937_qpcr_F | | tggagcctctcctccatttg |
| CDR20291_2937_qpcr_R | *CDR20291_2937* | cacatgcagctagtcaccct |
| gatB_qpcr_F | | atgctgttgcttttattgctgg |
| gatB_qpcr_R | *gatB* | atagctgaatgtgcaacaaagc |
| CDR20291_2938_qpcr_F | | atactgtggccactgctgtc |
| CDR20291_2938_qpcr_R | *CDR20291_2938* | actaggtgggacgacgtttg |
| CDR20291_1306_F | | ttagaagggaaaaatggggaagc |
| CDR20291_1306_R | *CDR20291_1306_F* | ttgaggaagtgtttttgcaact |
| thiD_qpcr_F | | acatggaacagggtgcacat |
| thiD_qpcr_R | *thiD* | actggtcctactccatgacct |
| CDR20291_1692_qpcr_F | | tggaagaccacaacctgcat |
| CDR20291_1692_qpcr_R | *CDR20291_1692* | ataggtggagttgctgctgg |
| CDR20291_1691_qpcr_F | | ttagggcatccaccaacacc |
| CDR20291_1691_qpcr_R | *CDR20291_1691* | acacaggcacttgctactga |
| CDR20291_3504_qpcr_F | | tgtttggcattgcatgagctg |
| CDR20291_3504_qpcr_R | *CDR20291_3504* | tggtccagctataatggggc |
| CDR20291_0781_qpcr_F | | gcccaacttgcatgaaaaagc |
| CDR20291_0781_qpcr_R | *hsmA* | tgagtaacatagccaacagcca |
| CDR20291_1308_nim_RT_F | | acttatgtccatttcttgcacaatg |
| CDR20291_1308_nim_RT_R | *nimB* | tggtgaatttggtactttttccact |
| hcp_qpcr_F | | tggaggttgtgatggtgcaa |
| hcp_qpcr_R | *hcp* | agccacctatatccccaaggt |
| recA_qpcr_F | | tgcaaaagctttgggtgtgg |
| recA_qpcr_R | *recA* | gcctttggaactaatgctgct |
| dnak_qpcr_F | | atcccctgctaaaactccagc |
| dnak_qpcr_R | *dnak* | accagcagtacaagaagctgtt |

| | | |
|---|---|---|
| 16rRNA_qpcr_F | | ctgggagacttgagtgcagg |
| 16rRNA_qpcr_R | *16rRNA* | gcctcagcgtcagttacagt |
| | **qPCR knockdown** | |
| nimB_qpcr_KD_F2 | *nimB* | tgtttaaagaaatgagattgaaaaaaagagaaatgacaaaaga |
| nim_qpcr_KD_R2 | | agcaactccataaggataaccattttcaga |
| hatT_qpcr_KD_F | | gtggtgtttaccttgaatcataatact |
| hatT_qpcr_KD_R | *hatT* | cagaagcgtatgttaatgtgtatatggt |
| hatR__qpcr_KD_F | | atgccaaagattttagaaaatgtaaaaga |
| hatR_qpcr_KD_R | *hatR* | tatatactgtacctgcacttataccacacctt |
| hsmA_qpcr_KD_F | | atgaattataaattaatacttgcaat |
| hsmA_qpcr_KD_R | *hsmA* | cctgtaatactatgaaggttttgtgata |
| hsmR_qpcr_KD_F | | gtgattttattgaaaagtaattatgactg |
| hsmR_qpcr_KD_R | *hsmR* | gcctgaaaccgtacctttagc |
| nimB_H61A_F | | cattgtgcaagaaatggagcaaagttagataatatatcg |
| nimB_H61A_R | | cgatatattatctaactttgctccatttcttgcacaatg |
| nimB_H55A_F | | aatgattctatttactttgcatgtgcaagaaatggacat |
| nimB_H55A_R | | atgtccatttcttgcacatgcaaagtaaatagaatcatt |
| nimB_H147A_F | | gttaaaatagaaattgacgcaatttctggtaaagcttca |
| nimB_H147A_R | | tgaagctttaccagaaattgcgtcaatttctattttaac |
| nimB_C101A_F | NimB site-directed mutagenesis | atagtttttggaaaagctgcaactgtagaaaatgaagaa |
| nimB_C101A_R | | ttcttcattttctacagttgcagcttttccaaaaactat |
| nimB_S67K_F | | cataagttagataatataaaaaaaaacaacaaagtttct |
| nimB_S67K_F | | agaaactttgttgtttttttttatattatctaacttatg |
| nimB_C56A_F | | gattctatttactttcatgcagcaagaaatggacataag |
| nimB_C56A_R | | cttatgtccatttcttgctgcatgaaagtaaatagaatc |
| nimB_Y119A_F | | gtagaaataataaaaaaagcatctaaaggattctttgaa |
| nimB_Y119A_F | | ttcaaagaatcctttagatgctttttttattatttctac |
| AV_sacI_F | | taagcagagctccctcgagtaaattcgaacacaacg |
| AV_BamHI_R | Anf3 cloning | tgcttaggatccttatcctaatctaaattttaat |
| BF_sacI_F | | taagcagagctcacgaagatccactattcgta |
| BF_BamHI_R | NimB_*B. fragilis* cloning | tgcttaggatccttatggtctatcatttctatttttagt |
| Inv_seq_F | | tccttcttaattacaaatttttagca |
| Inv_seq_R | inverse PCR | gagtcgcttttgtaaatttggaa |
| nimB_del_F | | ggatttactcaccctagagaag |
| nimB_del_R | nimB deletion confimation | cttgaagaaatgaatatcgaggta |

**Supplementary Table 3.** Genetically constructed strains from this study.

| Strains/contsructs | Note | Reference |
|---|---|---|
| R20291Tn::*nimB* | Tn mutant | This Study |
| R20291Tn::*cwp84* | | This Study |
| R20291Δ*nimB* | *nimB* knockout | This Study |
| R20291Δ*nimB* pRPF185 | empty vector pRPF185 | This Study |
| R20291 pRPF185 | | This Study |
| CD196 pRPF185 | | This Study |
| R20291Tn::*nimB* pRPF185 | | This Study |
| CD630 pRPF185 | | This Study |
| R20291Δ*nimB* pRPF185-P*nimB* $_{R20291}$ | *nimB* expression under *nimB* promoter | This Study |
| R20291 pRPF185-P*nimB* $_{R20291}$ | | This Study |
| R20291 pRPF185-P*nimB* $_{CD196}$ | | This Study |
| CD196 pRPF185-P*nimB* $_{R20291}$ | | This Study |
| CD196 pRPF185-P*nimB* $_{CD196}$ | | This Study |
| R20291Tn::*nimB* pRPF185-P*nimB* $_{R20291}$ | | This Study |
| R20291Tn::*nimB* pRPF185-P*nimB* $_{CD196}$ | | This Study |
| R20291 pRPF185-P$_{ATc}$*nimB* $_{R20291}$ | *nimB* expression under P*tet* promoter | This Study |
| R20291 pRPF185-P$_{ATc}$*nimB* $_{CD196}$ | | This Study |
| CD196 pRPF185-P$_{ATc}$*nimB* $_{R20291}$ | | This Study |
| CD196 pRPF185-P$_{ATc}$*nimB* $_{CD196}$ | | This Study |
| R20291Tn::*nimB* pRPF185-P$_{ATc}$*nimB* $_{R20291}$ | | This Study |
| R20291Tn::*nimB* pRPF185-P$_{ATc}$*nimB* $_{CD196}$ | | This Study |
| R20291 pMSPT | empty vector pMSPT | This Study |
| 70/76 pMSPT | | This Study |
| CD196 pMSPT | | This Study |
| JH7 pMSPT | | This Study |
| R20291 pMSPT-*nimB* i | Translational knockdown | This Study |
| 70/76 pMSPT-*nimB* i | | This Study |
| JH7 pMSPT-*nimB* i | | This Study |
| CD196 pMSPT-*nimB* i | | This Study |
| R20291 pMSPT-*ptsG* i | | This Study |
| R20291 pMSPT-*hatT* i | | This Study |
| R20291 pMSPT-*hatR* i | | This Study |
| R20291 pMSPT-*hsmA* i | | This Study |
| R20291 pMSPT-*hsmR* i | | This Study |
| R20291 pXWxyl-dcas9 | Transcriptional knockdown | This Study |
| 70/76 pXWxyl-dcas9 | | This Study |
| CD196 pXWxyl-dcas9 | | This Study |
| JH7 pXWxyl-dcas9 | | This Study |
| R20291 pXWxyl-dcas9-*nimB* i | | This Study |
| JH7 pXWxyl-dcas9-*nimB* i | | This Study |
| 70/76 pXWxyl-dcas9-*nimB* i | | This Study |

| | | |
|---|---|---|
| CD196 pXWxyl-dcas9-*nimB* i | | This Study |
| R20291 pXWxyl-dcas9-*vanRS* i | | This Study |
| R20291 pDSW1728 | mcherry empty vector pDSW1728 | This Study |
| CD196 pDSW1728 | | This Study |
| JH160 pDSW1728 | | This Study |
| 70/76 pDSW1728 | | This Study |
| R20291 pDSW1728-P*nimB* $^{G}$ | P*nimB* $^{G}$ mcherry OR P*nimB* $^{T}$ mcherry | This Study |
| R20291 pDSW1728-P*nimB* $^{T}$ | | This Study |
| CD196 pDSW1728-P*nimB* $^{G}$ | | This Study |
| CD196 pDSW1728-P*nimB* $^{T}$ | | This Study |
| JH160 pDSW1728-P*nimB* $^{G}$ | | This Study |
| JH160 pDSW1728-P*nimB* $^{T}$ | | This Study |
| 70/76 pDSW1728-P*nimB* $^{G}$ | | This Study |
| 70/76 pDSW1728-P*nimB* $^{T}$ | | This Study |
| R20291 pRPF185-$P_{ATc}$*anf3* | heterologus Nim expression | This Study |
| R20291 pRPF185-$P_{ATc}$*nimB* $_{BF}$ | | This Study |
| CD196 pRPF185-$P_{ATc}$*anf3* | | This Study |
| CD196 pRPF185-$P_{ATc}$*nimB* $_{BF}$ | | This Study |
| CD630 pRPF185-$P_{ATc}$*anf3* | | This Study |
| CD630 pRPF185-$P_{ATc}$*nimB* $_{BF}$ | | This Study |
| R20291Δ*nimB* pRPF185-$P_{ATc}$*anf3* | | This Study |
| R20291Δ*nimB* pRPF185-$P_{ATc}$*nimB* $_{BF}$ | | This Study |
| CD196 pRPF185-P*nimB* H55A | *nimB* mutagenesis | This Study |
| R20291Tn::*nimB* pRPF185-P*nimB* H55A | | This Study |
| R20291Tn::*nimB* pRPF185-P*nimB* C56A | | This Study |
| R20291Tn::*nimB* pRPF185-P*nimB* H61A | | This Study |
| R20291Tn::*nimB* pRPF185-P*nimB* Y119A | | This Study |
| R20291Δ*nimB* pRPF185-P*nimB* H55A | | This Study |
| *E. coli* BL21(DE3) pWL613a | Protein expression | This Study |
| *E. coli* BL21(DE3) pWL613a-NimB | | This Study |
| *E. coli* BL21(DE3) pWL613a-NimB H55A | | This Study |
| *E. coli* SD46 | conjugation | |
| *E. coli* CA434 | | |
